# Programmed ER fragmentation drives selective ER inheritance and degradation in budding yeast meiosis

**DOI:** 10.1101/2021.02.12.430990

**Authors:** George M. Otto, Tia Cheunkarndee, Jessica M. Leslie, Gloria A. Brar

## Abstract

The endoplasmic reticulum (ER) is a membrane-bound organelle with diverse, essential functions that rely on the maintenance of membrane shape and distribution within cells. ER structure and function are remodeled in response to changes in cellular demand, such as the presence of external stressors or the onset of cell differentiation, but mechanisms controlling ER remodeling during cell differentiation are not well understood. Here, we describe a series of developmentally regulated changes in ER morphology and composition during budding yeast meiosis, a conserved differentiation program that gives rise to gametes. During meiosis, the cortical ER undergoes fragmentation before collapsing away from the plasma membrane at anaphase II. This programmed collapse depends on the meiotic transcription factor Ndt80, conserved ER membrane structuring proteins Lnp1 and reticulons, and the actin cytoskeleton. A subset of ER is retained at the mother cell plasma membrane and excluded from gamete cells via the action of ER-plasma membrane tethering proteins. ER remodeling is coupled to ER degradation by selective autophagy, which is regulated by the developmentally timed expression of the autophagy receptor Atg40. Autophagy relies on ER collapse, as artificially targeting ER proteins to the cortically retained ER pool prevents their degradation. Thus, developmentally programmed changes in ER morphology determine the selective degradation or inheritance of ER subdomains by gametes.

## Introduction

The endoplasmic reticulum (ER) is a membrane-bound organelle that carries out a range of essential and conserved cellular functions, including protein synthesis and trafficking, lipid metabolism and inter-organelle communication. These functions rely on the maintenance of ER structure and subcellular distribution, which are achieved through membrane-shaping proteins, fusion and fission of ER tubules, and tethering between the ER and other cellular structures, including organelles and the plasma membrane (reviewed in Schwarz and Blower, 2016; Westrate et al., 2015). ER structure is highly dynamic even in unperturbed cells, and is dramatically remodeled in response to changes in cellular demand, such as protein folding stress or cell differentiation. Mutations that disrupt ER morphology are linked to a range of neurodegenerative diseases, including Alzheimer’s disease, amyotrophic lateral sclerosis, and hereditary spastic paraplegia (Öztürk et al., 2020; Renvoisé and Blackstone, 2010), highlighting the intimate connection between ER structure and function, and the importance of ER quality control during cell differentiation.

The ER emanates from the nuclear envelope and localizes around the nucleus (perinuclear ER) as well as the cell periphery (cortical ER) where it forms extensive contacts with the plasma membrane. In budding yeast, ER-plasma membrane (ER-PM) contacts are maintained by at least six tethering proteins, including Ist2, the tricalbins Tcb1, Tcb2 and Tcb3, and the vesicle-associated membrane protein-associated protein (VAP) orthologs Scs2 and Scs22 (Manford et al., 2012). All six tethers are integral ER membrane proteins that interact with phospholipids or proteins on the plasma membrane. Cells lacking these tethers have dramatically reduced cortical ER, disrupted lipid homeostasis, and acute sensitivity to ER stress, underscoring the importance of membrane tethering in maintaining ER structure and function. A second class of proteins involved in structuring the cortical ER is the reticulons, which form wedge-like structures in the cytosolic leaflet of the ER membrane to promote membrane curvature and drive the formation of ER tubules (Hu et al., 2008; Voeltz et al., 2006). ER tubules are highly dynamic, constantly growing, retracting, and fusing with one another to generate threeway tubule junctions (Guo et al., 2018). Fusion is mediated by the dynamin-like GTPases Sey1 (in budding yeast) or Atlastin (in metazoans) (Anwar et al., 2012; Hu et al., 2009; Orso et al., 2009). Lunapark (Lnp) family proteins are involved in the maintenance of three-way junctions and display functional antagonism with Sey1/Atlastin though the precise mechanistic role of Lnp in this process remains unclear (Chen et al., 2012, 2015; Wang et al., 2016). While factors that define ER structure are conserved across eukaryotes, we are only beginning to understand the diverse ways in which ER morphology and dynamics promote ER function.

Despite the fundamental relationship between ER structure and function, our knowledge of how the ER is remodeled as cells adapt to changing cellular conditions is limited. In budding yeast and cultured mammalian cells, exposure to chemical reducing agents causes ER protein folding stress and activation of the ER unfolded protein response (UPR), resulting in altered ER morphology and increased ER volume (Fumagalli et al., 2016; Schuck et al., 2009; Walter and Ron, 2011). Both ER stress and nutrient starvation drive selective degradation of the ER by autophagy (ERphagy), a response which is essential for cell adaptation and survival in these conditions (Fumagalli et al., 2016; Khaminets et al., 2015; Mochida et al., 2015; Zhang et al., 2020). While these studies provide crucial insight into ER quality control pathways that respond to cellular stress, the relationship between harsh drug treatment or prolonged starvation and the physiological conditions under which ER remodeling occurs is unclear. Here, we leverage the natural developmental context of budding yeast meiosis to study programmed ER remodeling in real time.

Meiosis is a conserved cell differentiation program that produces gamete cells specialized for sexual reproduction. In meiosis, a diploid progenitor cell undergoes a single chromosome duplication event followed by homolog pairing, recombination and two successive rounds of chromosome segregation, resulting in genetically distinct haploid gametes. In addition to ensuring the proper distribution of chromosomes, cells undergoing meiosis must deliver a full complement of cellular components into gametes while preventing the inheritance of toxic or deleterious material (Goodman et al., 2020; Neiman, 2011). While the regulation of meiotic chromosome segregation is well studied, mechanisms governing the inheritance and elimination of other cellular components during meiosis are relatively poorly defined.

In this study, we define key steps and mechanisms in ER inheritance and quality control in budding yeast meiosis. We find that during meiosis the cortical ER fragments and collapses away from the plasma membrane, a process which depends on the meiotic transcription factor Ndt80 but not chromosome segregation itself. Fragmentation and collapse rely on Lnp1, reticulon proteins and the actin cytoskeleton. A subset of ER fragments is retained at the plasma membrane and excluded from gametes in an ER-PM tether-dependent manner. In late meiosis, the ER is subject to extensive degradation by a selective autophagy mechanism that requires cortical ER collapse. Together, our work defines a developmental quality control mechanism in which programmed changes in ER morphology determine both the inheritance and selective exclusion of ER subdomains by gamete cells.

## Results

### The ER detaches from the plasma membrane during meiosis

Meiotic differentiation involves regulated partitioning of organelles to ensure the development of healthy spores. In order to characterize ER dynamics during meiotic differentiation, we used time-lapse microscopy to monitor cells expressing fluorescent markers of the ER lumen (GFP-HDEL) and chromatin (Htb1-mCherry). Premeiotic cells displayed ER morphology that is characteristic of mitotic cells, with ER distributed around the cell periphery (cortical ER) and the nucleus (perinuclear ER). As cells progressed through meiosis, the cortical ER underwent a striking series of morphological changes. Early in meiosis, just prior to the first nuclear division, the cortical ER coalesced into bright, highly dynamic rope-like structures, a phenomenon we refer to as “ER cabling” (figure 1A, S1A-B, video 1-2). Next, concurrent with anaphase II, the ER detached from the cell periphery and abruptly relocalized to an area in the center of cells roughly bounded by the four gamete nuclei (figure 1A-B, video 1-2). We refer to the abrupt detachment of cortical ER as “ER collapse”, a phenomenon that was previously predicted based on imaging of fixed cells in late meiotic stages, but that has not yet been studied in live cells (Suda et al., 2007). Finally, as spore packaging progressed, collapsed ER was inherited by each gamete and returned to the characteristic cortical and perinuclear structures seen in premeiotic cells (figure 1A, E).

**Figure 1.**
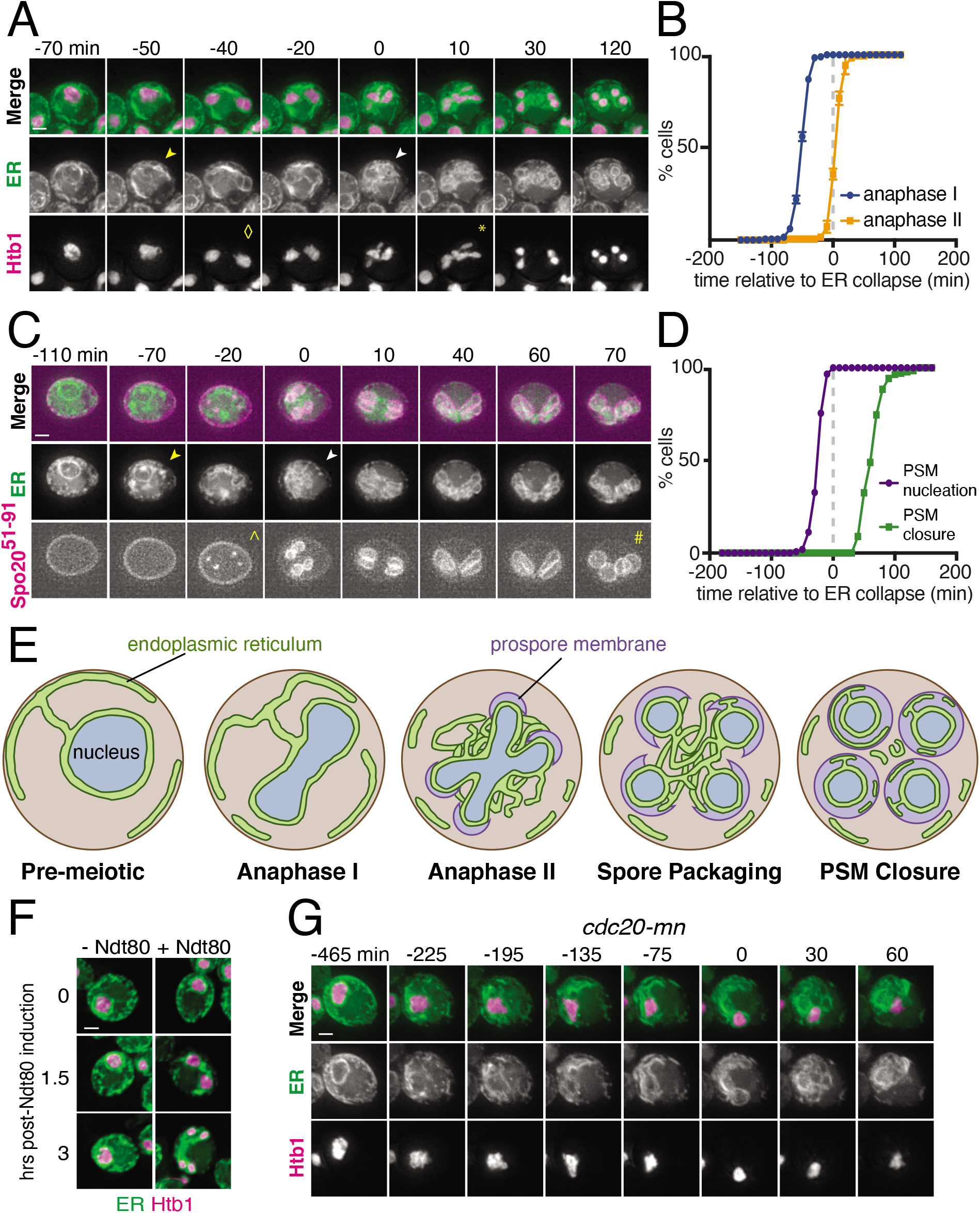
The ER undergoes developmentally regulated structural remodeling during meiosis. (A) Time-lapse microscopy of cells expressing GFP-HDEL to mark the ER (ER) and Htb1-mCherry to mark chromatin (Htb1) imaged every 10 minutes during meiosis. Symbols mark the onset of ER cabling (yellow arrowhead), ER collapse (white arrowhead), anaphase I (◇) and anaphase II (*). ER collapse is defined to occur at 0 min. (B) Quantification of the time of anaphase I and anaphase II relative to ER collapse. (C) Time-lapse microscopy of cells expressing GFP-HDEL (ER) and mKate-Spo20^51-91^ to mark the prospore membrane (PSM). Symbols mark the onset of ER cabling (yellow arrowhead), ER collapse (white arrowhead), PSM nucleation (^) and PSM closure (#). (D) Quantification of the time of PSM nucleation and closure relative to ER collapse. (E) Schematic of meiosis-coupled ER remodeling with organelles and stages of meiosis and spore formation labeled. (F) Cells expressing GFP-HDEL (ER), Htb1-mCherry (Htb1) and an estrogen-inducible allele of *NDT80* treated with 1 μM β-estradiol (+ Ndt80) or vehicle (-Ndt80) after 5 hours in sporulation media and imaged at the indicated times following induction. (G) As in (A) but in cells with the endogenous promoter of *CDC20* replaced with the mitosis-specific *CLB2* promoter (*cdc20-mn*), and cells imaged every 15 minutes. Scale bar = 2 μm for all panels.

In budding yeast, meiosis is coupled to spore formation, in which gamete plasma membranes (also called prospore membranes) are synthesized de novo and grow to encapsulate the full complement of cellular material to be inherited by gametes (Neiman, 2011). Imaging the ER alongside the marker of prospore membrane synthesis mKate-Spo20^51-91^ (Nakanishi et al., 2004) revealed that ER collapse takes place after prospore membrane nucleation but prior to closure (figure 1C-D, video 3). Based on the timing of ER collapse and the spatial relationship between collapsed ER and nascent prospore membranes, it appears that cortical ER collapse is needed for its delivery into gamete cells.

The precise timing with which ER detachment takes place relative to meiotic chromosome segregation and prospore membrane formation suggests that this process is tightly regulated as part of the broader developmental program that coordinates meiosis and spore formation. To further test this idea, we disrupted meiotic progression and assessed the impact on ER dynamics. First, we arrested cells in prophase I by withholding the meiotic transcription factor Ndt80, which is required to initiate the two meiotic nuclear divisions following homologous recombination (Benjamin et al., 2003; Chu and Herskowitz, 1998). Arrested cells did not undergo ER cabling or collapse, indicating that these processes depend on Ndt80 induction and are not simply a response to the nutrient-poor conditions that stimulate meiosis in budding yeast. (figure 1F Blocking meiotic chromosome segregation using a meiotic null allele of the anaphase-promoting complex/cyclosome (APC/C) activator Cdc20 (*cdc20-mn*; Lee and Amon, 2003), however, did not prevent. cortical ER detachment from the plasma membrane. Cortical ER coalesced around the single, undivided nucleus in *cdc20-mn* cells similar to what occurs around anaphase II in wild type cells (figure 1H). (figure 1G, video 4). Together, these data indicate that meiotic ER remodeling is triggered by a developmental cue downstream of Ndt80, but is independent of chromosome segregation and the consequent dramatic changes to nuclear morphology.

### ER-plasma membrane tethers define a cortically retained ER compartment

How is the abrupt detachment of the ER from the plasma membrane (PM) achieved? In budding yeast, at least six proteins function as ER-PM tethers. These include Ist2, the tricalbins Tcb1, Tcb2 and Tcb3, and the VAP orthologs Scs2 and Scs22. Cells lacking all six tethers have drastically reduced levels of cortical ER, disrupted lipid homeostasis, and reduced tolerance to ER stress (Manford et al., 2012). We sought to determine the role of ER-PM tethering proteins in meiotic ER collapse by imaging each tether during meiosis. To our surprise, Ist2 and all three tricalbins remained cortically localized throughout meiosis, even during anaphase II when the ER has collapsed (figure 2A, S2A-D, video 5-8). In contrast, Scs2 and Scs22 demonstrated collapsed morphology at anaphase II (figure 2A, S2E-F, video 9-10).

**Figure 2.**
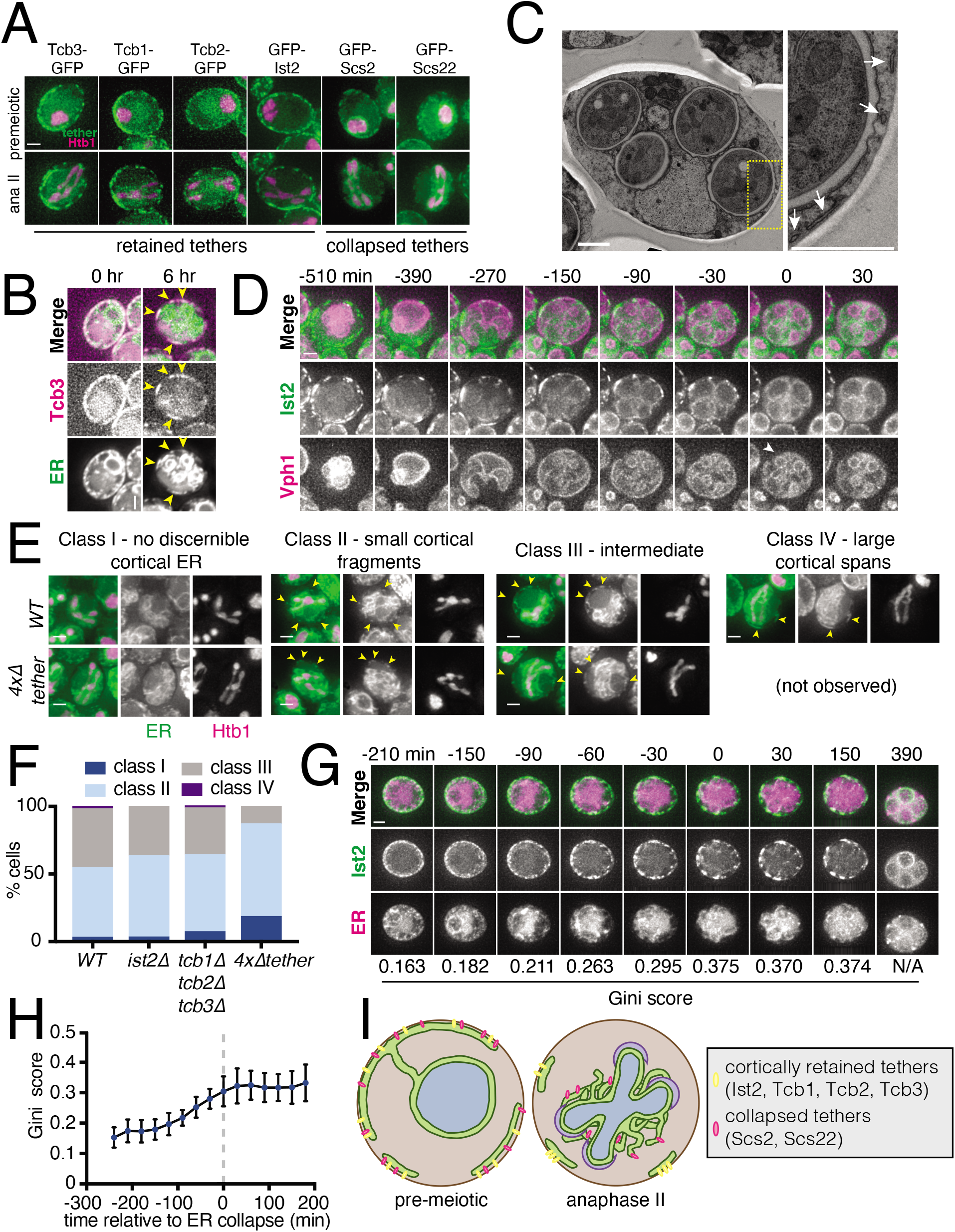
A subset of ER-PM tethering proteins marks cortically retained ER islands. (A) Time-lapse microscopy of cells expressing the indicated ER-PM tether tagged with GFP (tether) and Htb1-mCherry (Htb1) imaged during meiosis. A representative cell is shown prior to meiosis (top) and during anaphase II (bottom). Tethers are categorized as retained or collapsed based on anaphase II localization. (B) Cells expressing GFP-HDEL (ER) and Tcb3-mKate (Tcb3) imaged at 0 hr (top) and 6 hr (bottom) after introduction to sporulation media. Arrowheads mark islands of colocalized ER lumen and Tcb3 signal. (C) Electron microscopy of a WT cell following spore closure (left). Yellow box outlines the area shown zoomed-in on the right. White arrows indicate cortically retained ER fragments. “S” marks the four spores. “V” marks the vacuole. (D) Time-lapse microscopy of cells expressing GFP-Ist2 and Vph1-mCherry to mark the vacuole imaged every 30 minutes in meiosis. Arrowhead marks the time of vacuole lysis, which is defined as time 0. (E) Images of WT or *ist2Δ tcb1Δ tcb2Δ tcb3Δ (4xΔtether*) cells expressing GFP-HDEL (ER) and Htb1-mCherry (Htb1) taken at anaphase II. When possible, a representative cell of each genotype is shown for each indicated cortical ER classification. (F) Quantification of at least 100 cells for the indicated genotypes following the classification system in (E). (G) Time-lapse microscopy of cells expressing GFP-Ist2 and mCherry-HDEL (ER) imaged every 30 minutes in meiosis. The Gini score based on quantification of Ist2 signal is shown below each timepoint. Minute 0 is defined as the time of ER collapse. (H) Gini quantification based on cortical Ist2 signal over time. Values are the average of 10 cells scored across each timepoint. Error bars represent standard deviation. (I) Schematic showing ER morphology and tether localization in pre-meiotic and anaphase II cells. Scale bar = 2 μm for all panels except C, for which scale bar = 1 μm.

The cortical retention of a subset of ER-PM tethers was unexpected because all four proteins have integral membrane domains anchoring them in the ER and are therefore predicted to localize with the ER. We therefore wondered if the cortically retained ER-PM tethers represented previously overlooked fragments of ER that failed to detach from the PM during ER collapse. Imaging Tcb3-mKate alongside GFP-HDEL revealed that Tcb3 signal at the cell cortex indeed overlapped with small islands of ER lumen, even when the vast majority of the ER was collapsed (figure 2B). Analysis of previously published electron microscopy of meiotic cells revealed small fragments of ER that remained bound to the plasma membrane even after prospore membrane closure (figure 2C; King et al., 2019). Together, our observations indicate that a subset of ER-PM tethers define a previously unappreciated cortically retained ER compartment.

Because the gamete plasma membrane is formed de novo rather than inherited from the progenitor cell, any cellular component that is attached to the progenitor plasma membrane is necessarily excluded from gametes. We observed an abrupt decrease in the signal of all four excluded tethers in late meiosis, suggesting that excluded ER fragments are degraded during this time (video 5-8). Late in meiosis, the yeast vacuole dramatically expands before ultimately lysing, releasing its contents into the ascoplasm region outside of spores and degrading the excluded material, including protein aggregates and nuclear pore complexes (Eastwood et al., 2012; King et al., 2019). To see if this is also the mechanism responsible for eliminating cortically retained ER, we performed time-lapse imaging of cells expressing Vph1-mCherry along with Tcb3-GFP or GFP-Ist2. Vacuole lysis, indicated by a switch in mCherry signal from vacuole membrane-localized to diffuse, coincided perfectly with the disappearance of cortical ER signal, supporting a model in which the release of vacuolar proteases into the ascoplasm is responsible for the degradation of cortically retained ER (figure 2D, S2G, video 11-12). Thus, cortical ER retention is a means by which cells can exclude and degrade certain parts of the ER from gamete cells.

### ER-PM tethers promote the cortical retention of ER fragments during ER collapse

Cortically retained ER fragments appear to represent a small portion of the cell’s total ER pool. Nonetheless, we observed cell-to-cell heterogeneity in the amount of retained cortical ER in our live-cell microcopy experiments. To quantify this heterogeneity, and to enable us to assess the effect of genetic manipulation on cortical ER retention, we classified cells into four distinct groups: class I (no discernable ER retention), class II (small cortical ER fragments), class III (intermediate between classes II and IV), and class IV (large spans of cortical ER) (figure 2E). The vast majority of wild-type cells scored at anaphase II of meiosis fell into classes II and III, though we did observe a small number of class I and class IV cells (figure 2F). Deletion of the four cortically retained tethers (4xΔtether) resulted in a significant increase in the frequency of cells falling into class I and fewer cells in class III, indicating an overall decrease in the amount of retained cortical ER in this genotype (figure 2F, S2J, video 13). Cells lacking only a subset of these four tethers showed an intermediate phenotype, suggesting that the loss of ER-PM tethers has an additive effect on the amount of retained cortical ER (figure 2F, S2H-I, video 14-15). We conclude that Ist2 and the tricalbins promote the exclusion and subsequent degradation of cortical ER fragments in meiosis.

### Clustering of ER-PM tethers precedes ER collapse

We noted that although the cortically retained ER-PM tethers did not undergo collapse with the bulk of the ER, their localization was not static over time. Early in meiosis, tether signal was distributed somewhat homogenously around the cell cortex. However, this pattern changed as meiosis progressed, with tether signal becoming more clustered over time, resulting in tether-rich islands separated by stretches of cell cortex with no tether signal (figure 2G, video 16). To quantitatively assess the degree to which tether signal is asymmetrically distributed within the cell cortex, we employed a metric called the Gini coefficient (G), which measures inequality within a dataset on a scale from zero to one (Rouskin et al., 2014; Wittebolle et al., 2009). If tether signal were distributed perfectly evenly throughout the cell cortex, it would receive a Gini coefficient of zero, whereas a highly asymmetric distribution of signal would be closer to one. Cells in early meiosis had a relatively low Gini coefficient for GFP-Ist2 distribution (G = 0.153 +/- 0.034) (figure 2G-H). This value steadily increased over time before plateauing (G = 0.304 +/- 0.049) at the time of ER collapse. This analysis demonstrates that the onset of tether clustering precedes ER collapse by several hours and therefore represents an early step in meiotic ER remodeling. Though the exact relationship between tether clustering and ER collapse is still unclear, our observations support a model in which the cortical ER is sorted into tether-containing (cortically retained) and tether-free (collapsed) domains to allow selective ER retention and inheritance (figure 2I).

### Reticulons promote ER detachment

How is the normally continuous cortical ER separated into collapsed and retained pools? One means of separating a continuous compartment into two separate topologies is by membrane fission, a phenomenon underlying key biological processes such as endocytosis, mitochondrial division, and cytokinesis. While the molecular mechanisms driving many membrane fission events are well characterized, the regulation of membrane fission in the ER is relatively poorly defined. Nevertheless, a growing body of evidence supports a role for membrane curvature in driving ER tubule fission *in vitro* and *in vivo*. ER membrane curvature is regulated by reticulons, a conserved class of proteins that generate ER membrane curvature via a double-hairpin reticulon homology domain (RHD) (Hu et al., 2008; Voeltz et al., 2006). Overexpression of reticulon proteins results in ER fragmentation in cell culture and Drosophila models, while *in vitro* reconstituted ER networks containing reticulons spontaneously fragment in the absence of fusion-promoting factors (Espadas et al., 2019; Powers et al., 2017; Wang et al., 2016). These observations led us to pursue the hypothesis that reticulons mediate the topological separation of retained and collapsed ER by promoting membrane fission.

Budding yeast have two reticulons, Rtn1 and Rtn2, as well as a reticulon-like protein Yop1, that together are required for normal ER tubule formation (Voeltz et al., 2006). As expected, *rtn1Δ rtn2Δ yop1Δ* mutants displayed a drastic reduction in tubules and an increase in ER sheet structures under mitotic growth conditions (figure S3A). Additionally, in meiotic cells, we observed a striking increase in the amount of ER that remained cortically localized beyond anaphase II (figure 3A-B, video 17). Relative to WT, *rtn1Δ rtn2Δ yop1Δ* cells were much less likely to have class II cortical ER at anaphase II and much more likely to fall into class IV. Moreover, the cabling behavior that we observe in WT cells immediately prior to collapse was absent in *rtn1Δ rtn2Δ yop1Δ* cells, suggesting that membrane curvature and/or fission are important for ER cabling, and that the cabling process may promote cortical ER detachment (figure 3A, S3B video 18). These observations support a role for reticulon-mediated membrane curvature in promoting meiotic ER collapse via fragmentation.

**Figure 3.**
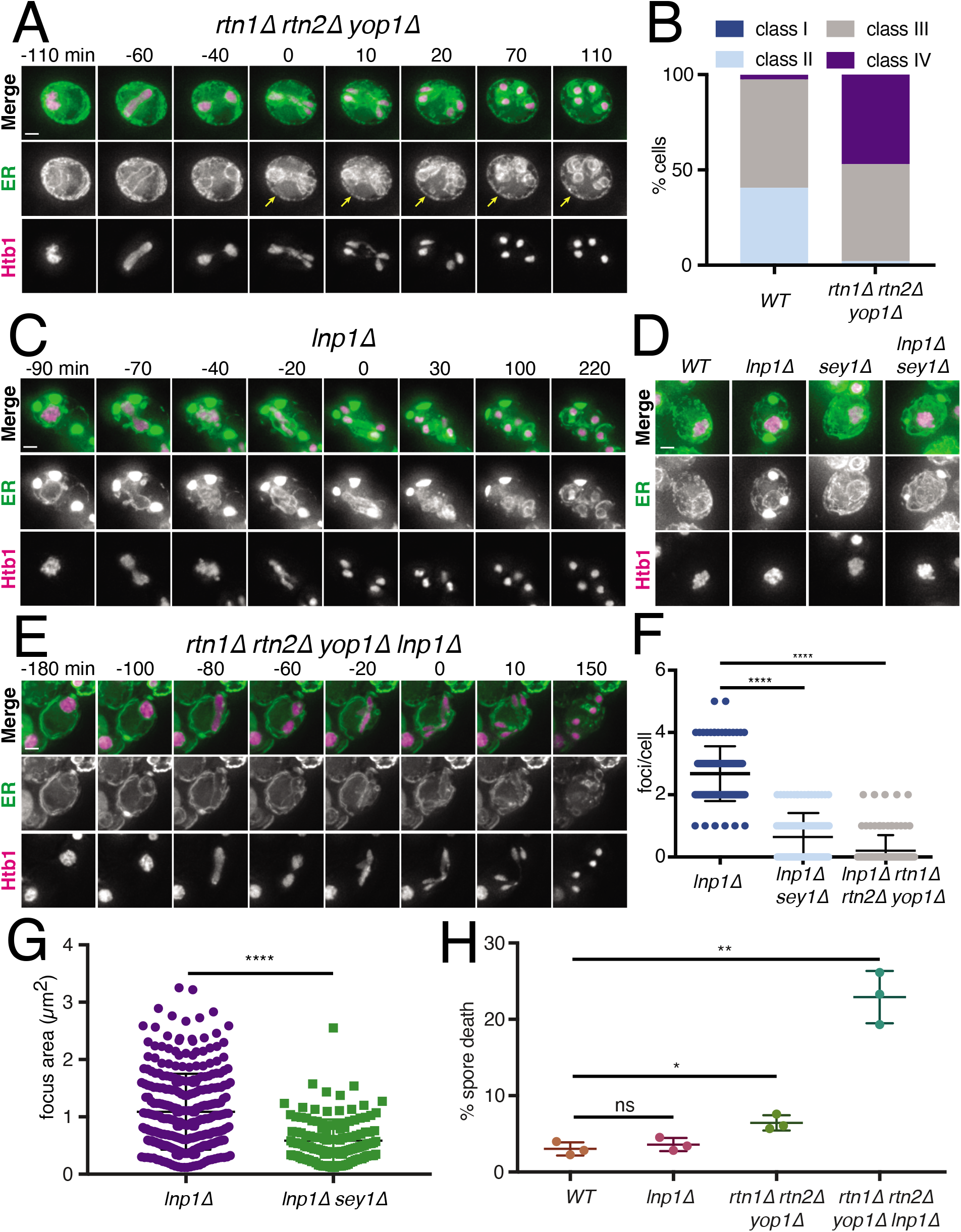
Reticulons and Lnp1 regulate meiotic ER remodeling. (A) Time-lapse microscopy of rtn1Δ rtn2Δ yop1Δ cells expressing GFP-HDEL (ER) and Htb1-mCherry imaged every 10 minutes during meiosis. Minute 0 is defined as the time of ER collapse. Yellow arrows indicate abundant cortically retained ER persisting after anaphase II. (B) Classification of cortical ER retention for at least 100 cells for the indicated genotypes. (C) As in (A) but with cells of genotype lnp1Δ. (D) Cells of the indicated genotypes expressing GFP-HDEL and Htb1-mCherry imaged at 10 min in sporulation media. (E) As in (A) but with cells of genotype rtn1Δ rtn2Δ yop1Δ lnp1Δ. (F) Quantification of the number of foci per cell showing the average and standard deviation for of least 100 cells of the indicated genotypes. (G) Quantification of focus size for at least 100 cells of each of the indicated genotypes. Average and standard deviation are shown. (H) Spore viability quantification after 24 hr in sporulation media followed by germination for 48 hours on YPD. Each replicate represents results for at least 176 individual spores. Scale bar = 2 μm for all panels.

### Lnp1 is required for ER detachment

Both normal ER tubule fission in unperturbed cells and ER fragmentation upon reticulon overexpression can be countered by homotypic membrane fusion, which is carried out by the dynamin-like GTPase Sey1 (Atlastin in plants and metazoans) (Anwar et al., 2012; Espadas et al., 2019; Hu et al., 2009; Orso et al., 2009; Wang et al., 2016). While the factors regulating Sey1/Atlastin activity are incompletely defined, the lunapark protein Lnp1 displays functional antagonism with Sey1 in mitotic yeast cells (Chen et al., 2012). We reasoned that Lnp1 may promote ER collapse by negatively regulating Sey1-mediated ER membrane fusion. If this were the case, cells lacking Lnp1 would be expected to show increased ER retention.

To our surprise, and in contrast to mitotic cells, *lnp1Δ* mutants displayed massive cortical ER puncta when placed in sporulation media (figure 3C, S3C). We examined multiple ER markers, including luminal and transmembrane proteins, and found that all of them localized to large ER puncta in *lnp1Δ* cells, indicating that these aberrant structures are generally representative of ER in this condition (figure S3D). Consistent with a role for Lnp1 in promoting ER detachment, ER puncta in *lnp1Δ* cells retained their cortical localization throughout meiosis and spore packaging, resulting in their exclusion from gamete cells (figure 3C, video 19). Puncta in *sey1Δ lnp1Δ* double mutants were smaller and less abundant than those found in *lnp1Δ* mutants, suggesting that these phenotypes result from excessive Sey1-mediated membrane fusion (figure 3D, F-G).

To determine the relationship between reticulons and Lnp1 in promoting ER collapse, we examined ER dynamics in the quadruple *lnp1Δ rtn1Δ rtn2Δ yop1Δ* mutant. Strikingly, these cells rarely formed ER foci, suggesting that foci are normally comprised of highly reticulated ER (figure 3F). Moreover, they showed a dramatic increase in cortical ER retention during anaphase II, with all observed cells observed falling into class IV (figure 3E, video 20). We also found that *lnp1Δ rtn1Δ rtn2Δ yop1Δ* mutant cells had dramatically reduced spore viability, whereas mutants lacking only the three reticulons had a modest viability defect and *lnp1Δ* cells were unaffected (figure 3H). Together, these data reveal a role for the regulation of membrane shape and fusion in ensuring normal ER detachment during meiosis and, ultimately, the health of the gametes produced during this process.

### Artificial ER-PM tethering does not prevent ER collapse

Impaired ER collapse in cells lacking reticulons could result directly from reduced reticulon-dependent tubule severing, or indirectly from altered ER morphology. We thus asked whether we could artificially tether cortical ER to the plasma membrane throughout meiosis without altering reticulon levels (figure 4A). We constitutively tethered GFP-Scs2 to the cell cortex using the plasma membrane protein Pil1 fused to a genomically encoded antibody against GFP (Pil1-antiGFP; Schmit et al., 2018). Whereas GFP-Scs2 in WT cells localized with collapsed ER in anaphase II, GFP signal remained strictly cortical in cells expressing Pil1-antiGFP, indicating that our artificial tethering strategy was successful (figure 4B-C, video 21-22). To our surprise, this manipulation did not have a strong effect on overall cortical ER retention at anaphase II, as assessed by mCherry-HDEL localization, which was largely collapsed in late meiosis. This result shows that that introducing an artificial constitutive ER-PM tether does not prevent collapse, and, incidentally, that cortical release of the ER-PM tether Scs2 does not drive meiotic ER collapse.

**Figure 4.**
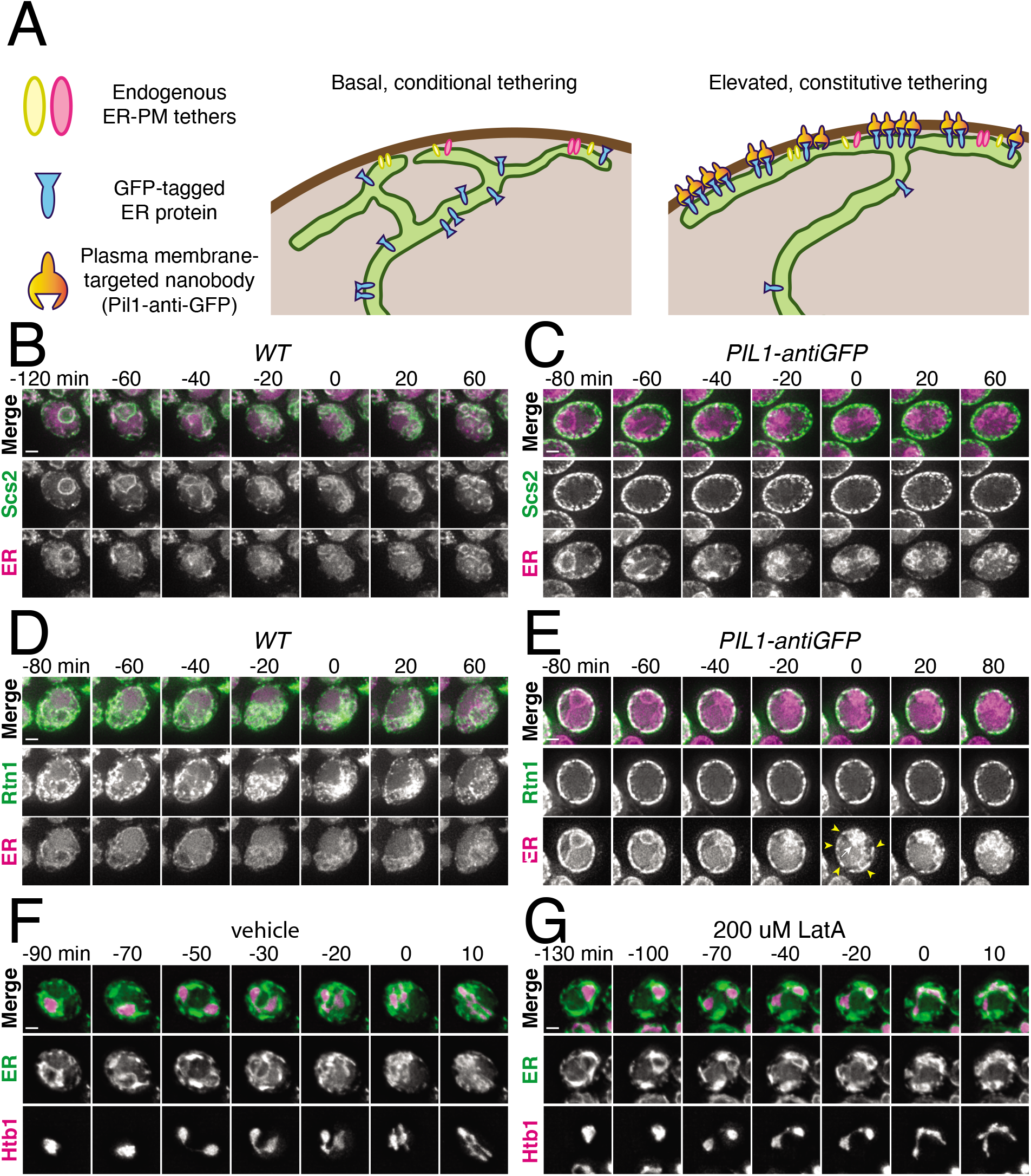
Artificial cortical ER tethering does not prevent ER collapse. (A) Schematic of artificial cortical ER tethering using Pil1-antiGFP. (B) Time-lapse microscopy of cells expressing GFP-Scs2 and mCherry-HDEL (ER) imaged every 10 minutes during meiosis. 0 min is defined as the time of ER collapse. (C) As in (B) but with cells expressing Pil1-antiGFP nanobody. (D) As in (B) but with cells expressing Rtn1-GFP instead of GFP-Scs2. Yellow arrowheads indicate cell cortex devoid of ER. White arrow indicates collapsed ER. (E) As in (D) but with cells expressing Pil1-antiGFP nanobody. (F) Cells expressing GFP-HDEL (ER) and Htb1-mCherry treated with DMSO (vehicle) at 4.5 hr in meiosis and imaged every 10 min. 0 min is defined as the onset of anaphase II. (G) As in (F) but cells were treated with 200 uM LatA instead of vehicle. Scale bar = 2 μm for all panels

We reasoned that forced tethering of a more abundant cortical ER protein may be necessary to prevent collapse, leading us to perform a similar approach to that described above, this time using the abundant reticulon protein Rtn1. Although forced tethering of Rtn1-GFP increased the overall amount of cortically retained ER, collapse was not prevented, as assessed by a substantial amount of mCherry-HDEL signal in the collapsed ER pool at anaphase II and a corresponding reduction in cortical mCherry-HDEL signal (figure 4D-E, video 23-24). These results support a model in which the cortical ER is prone to reticulon-dependent fragmentation during meiosis II, resulting in either collapse or cortical retention based on each ER fragment’s local association (or lack of association) with ER-PM tethers. Thus, introduction of an abundant artificial tether increases the amount of ER that is cortically retained but cannot prevent bulk ER dissociation from the plasma membrane.

### The actin cytoskeleton promotes ER collapse

The abrupt, coordinated movement of cortical ER away from the plasma membrane suggests the involvement of a force-generating mechanism rather than passive diffusion. During mitosis in yeast, ER tubules are delivered into the daughter cell along actin cables (Estrada et al., 2003). To determine whether the actin cytoskeleton is also involved in meiotic ER dynamics, we treated cells undergoing meiosis with Latrunculin A (LatA), a drug that prevents actin polymerization. LatA-treated cells were still able to undergo ER cabling, but cabled structures failed to collapse, instead remaining cortical throughout chromosome segregation, suggesting that cabled ER is pulled away from the plasma membrane along actin filaments (figure 4F-G, video 25-26).

### The ER undergoes turnover during meiosis

To determine if morphological ER remodeling in meiosis is accompanied by a fundamental change in the molecular composition of the ER, we analyzed the expression of all ER proteins over the course of meiosis in a previously published dataset with global matched measurements for protein abundance and translation levels (Brar et al., 2012; Cheng et al., 2018). Because cell number remains constant throughout meiosis, decreases in protein abundance over time indicate active degradation. We observed evidence for degradation for almost every ER-localized protein during mid- to late-meiosis in multiple waves, occurring concomitant with or after ER collapse. Of the 658 proteins characterized for ER localization, we also observed robust synthesis (above 50 RPKM) for 81.3% of them, with this synthesis occurring late in meiosis (after ER collapse) for 85.2% of this set (figure 5A, S4). Together, these data suggest large-scale turnover and resynthesis of many ER components during the meiotic program.

**Figure 5.**
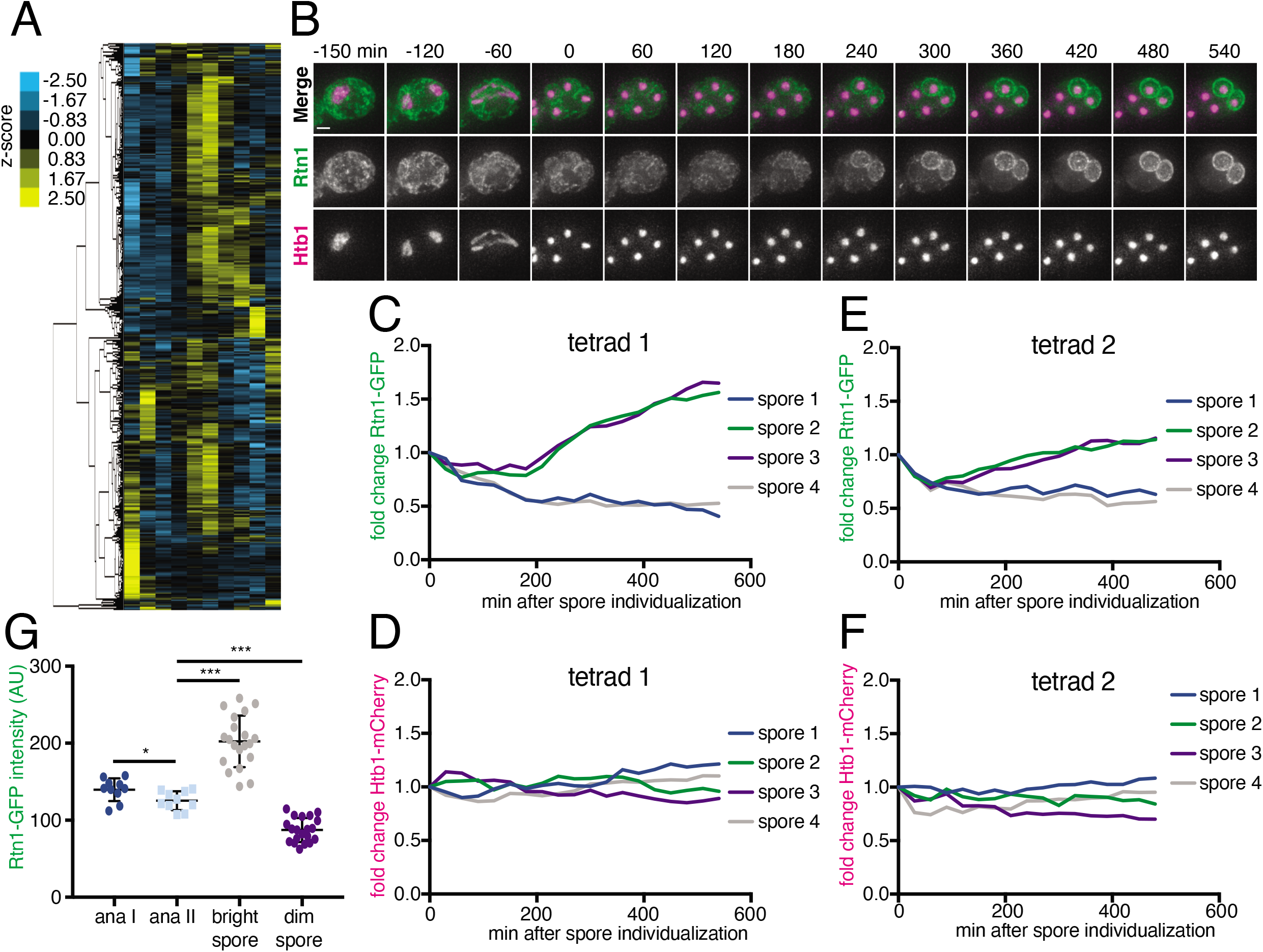
The ER undergoes turnover and resynthesis during meiosis. (A) Hierarchical clustering of z-score quantification of protein measurements published in Cheng *et. al*. (2018) for all quantified proteins annotated for ER localization. Each row represents one protein and each column is a timepoint in meiosis. (B) Time-lapse microscopy of cells heterozygous for *RTN1-GFP* and *HTB1-mCherry* imaged every 30 minutes in meiosis. Time 0 is defined as the time of spore individualization. Scale bar = 2 μm. (C) Quantification of the average GFP signal for each spore for the cell in (B). (D) Quantification of the average mCherry signal for each spore for the cell in (B). (E and F) as in (C) and (D) for a second representative cell. (G) Quantification of the average GFP signal for ten cells at anaphase I, anaphase II, and the last imaged timepoint separated based on signal brightness (bright spore and dim spore).

To independently assess the turnover of an abundant ER protein during meiosis, we used an assay that takes advantage of the diploid status of meiotic cells (Eisenberg et al., 2018) by imaging cells with heterozygous tags marking the ER (*RTN1-GFP/RTN1^WT^*) and histones (*HTB1-mCherry/HTB1^WT^*). In this system, preexisting ER is marked by Rtn1-GFP, whereas newly synthesized ER following spore closure will either be marked or unmarked, depending on if the spore inherited *RTN1-GFP* or *RTN1^WT^*, respectively. Similar GFP levels across all four spores would suggest no degradation or synthesis of Rtn1. A GFP signal increase in two spores would indicate Rtn1 resynthesis, while a GFP signal decrease in two spores indicates Rtn1 degradation. Consistent with our analysis of ER proteins globally, we saw two spores per tetrad retaining high levels of GFP signal, while GFP levels in the other two spores progressively declined, as opposed to the persistence of mCherry signal (and thus histone presence) in all four spores. This pattern provides further support for the existence of concerted ER degradation and resynthesis in late meiosis (figure 5B-G, video 27).

### The ER is degraded by autophagy during meiosis

What mechanisms mediate ER turnover during meiosis? One possibility was autophagy, in which cargo such as organelle fragments are engulfed by a double-membrane autophagosome and targeted to the vacuole (lysosome in metazoans) for degradation (Morishita and Mizushima, 2019). General autophagy factors are highly upregulated during meiosis and the kinase Atg1 is essential for both autophagy and entry into meiosis (Brar et al., 2012; Wen et al., 2016). Because GFP is resistant to vacuolar proteases while cargo proteins are not, GFP-tagged proteins that have been degraded by autophagy leave behind a GFP epitope that can be readily detected by western blot (Mochida et al., 2015). We tagged several ER resident proteins with GFP and observed the accumulation of a GFP-only band in meiosis, suggesting that the ER as a whole is a target of autophagy during this process (figure 6A, S5A-C). A faint GFP fragment was visible as early as a few hours into meiosis, but the greatest accumulation occurred as cells progressed through anaphase II and beyond (figure 6A, S5D). As an orthogonal means of observing ER autophagy (ERphagy), we imaged cells expressing Rtn1-GFP and Vph1-mCherry, a marker of the vacuole membrane. Prior to meiosis, there was very little GFP within the vacuole, whereas cells in late meiosis displayed strong, diffuse GFP signal throughout the vacuole, providing further evidence that cells induce ERphagy as they progress through meiosis (figure 6B).

**Figure 6.**
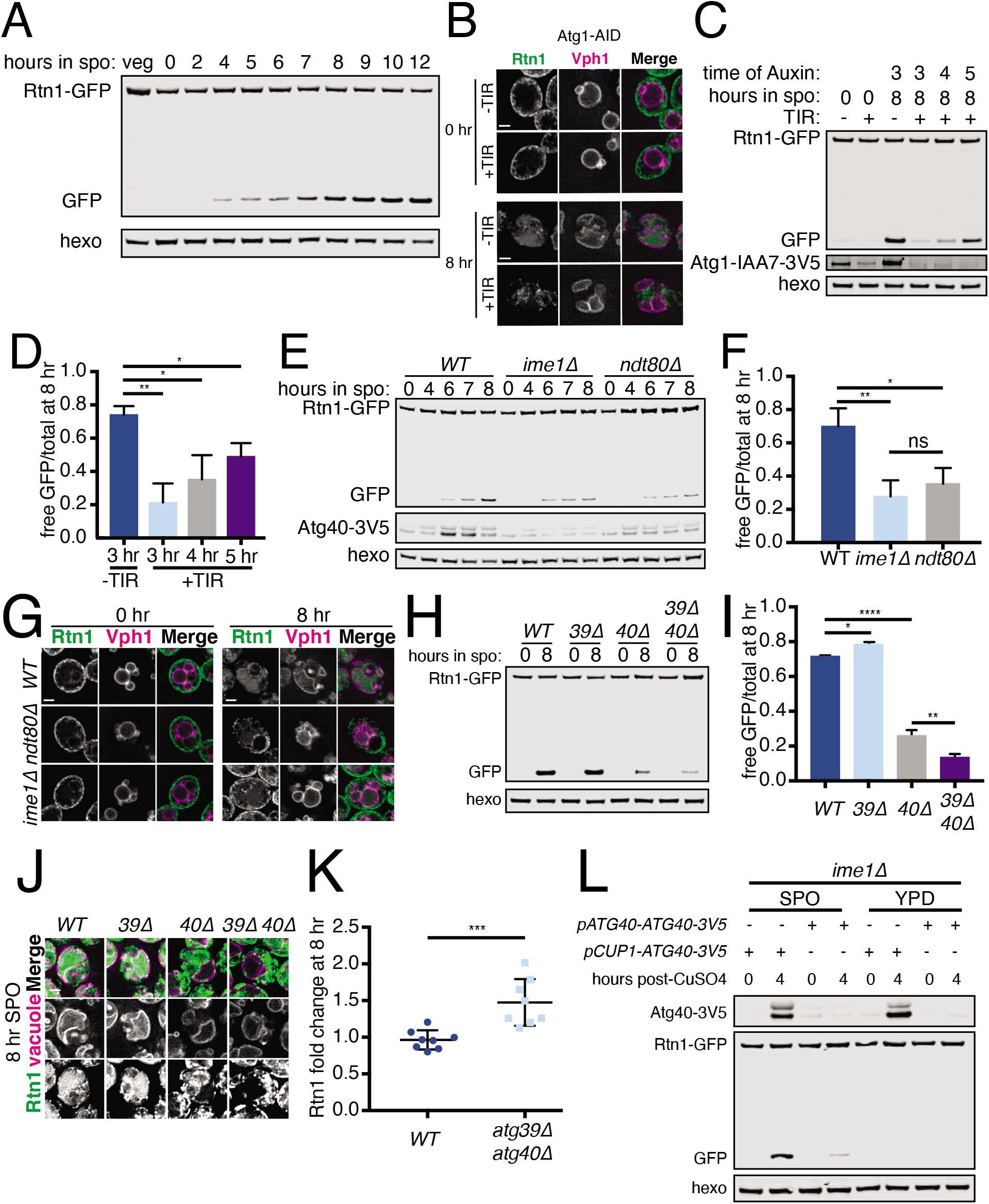
The ER is degraded by selective autophagy during meiosis. (A) Western blot with samples taken from cells expressing Rtn1-GFP during vegetative exponential growth (veg) or at the indicated time in meiosis probing for GFP and hexokinase (hexo) loading control. (B) Microscopy of cells expressing Rtn1-GFP, Vph1-mCherry, and Atg1-AID and imaged at the indicated times after transfer to sporulation media. Presence (+) or absence (-) of osTIR is indicated. Cells were treated with 500 μM auxin after 4 hr in sporulation media. (C) Western blot of cells of the same genotypes as in (B) probing for GFP, V5 and hexo. Cells were treated with 500 μM auxin at the indicated times. (D) Quantification of free GFP as a proportion of total GFP signal for three replicates of the experiment in (C). (E) Western blot with samples taken from cells of the indicated genotypes at the indicated times after transfer to sporulation media, probed for GFP, V5 and hexo. (F) Quantification of free GFP as a proportion of total GFP signal for three replicates of the experiment in (E). (G) Microscopy of cells of the indicated genotypes expressing Rtn1-GFP and Vph1-mCherry and imaged at the indicated times after transfer to sporulation media. (H) Western blot with samples taken from cells of the indicated genotypes expressing at the indicated times after transfer to sporulation media, probed for GFP and hexo. (I) Quantification of free GFP as a proportion of total GFP signal for three replicates of the experiment in (H). (J) Microscopy of cells of the indicated genotypes expressing Rtn1-GFP and Vph1-mCherry imaged at the indicated times following transfer to sporulation media. (K) Quantification of Rtn1-GFP signal at 8 hr following transfer to SPO normalized to hexokinase loading control and to 0 hr measurement for eight replicates of the experiment in (H). (L) Western blot with samples taken from *ime1Δ* cells expressing Rtn1-GFP and Atg40-3V5 under the endogenous promoter (*pATG40-ATG40-3V5*) or the *CUP1* promoter (*pCUP1-ATG40-3V5*). For YPD samples, cells were diluted to 0.05 ODU in YPD, allowed to grow to exponential phase, treated with50 μM CuSO_4_. For SPO samples, cells were transferred to sporulation media for 2 hours and treated with 50 μM CuSO_4_. Protein samples were taken at the indicated times after CuSO_4_ treatment. Scale bar = 2 μm for all panels.

Atg1 is required for entry into meiosis, so we could not assess the role of the canonical Atg1-dependent autophagy pathway in meiotic ERphagy using *atg1Δ* cells. Instead, we conditionally depleted cells of Atg1 after meiotic entry using the auxin-inducible degron system, in which TIR1, a plant-derived ubiquitin ligase, targets degron-bearing substrates for proteasomal degradation only in the presence of the plant hormone auxin (Nishimura et al., 2009). By withholding auxin until after meiotic entry, we were able to deplete cells of degron-tagged Atg1 (Atg1-AID) without blocking meiosis. Both the accumulation of Rtn1-GFP in the vacuole as seen by microscopy and the appearance of GFP as a lone fragment by western blot in Rtn1-GFP cells were strongly reduced by depletion of Atg1, an effect that was stronger the earlier cells were treated with auxin (figure 6B-D). Thus, ERphagy in meiosis takes place through the canonical Atg1-dependent pathway.

We next sought to determine whether ERphagy is induced as part of the developmental program of meiosis or simply in response to sporulation media, which is nutrient-poor. Cells progressing through meiosis induced ERphagy much more strongly than cells arrested in prophase I (*ndt80Δ*) or prior to meiotic entry (*ime1Δ*), indicating that this process is enhanced by meiotic progression (figure 6E-G). Interestingly, ERphagy differs from other forms of autophagy in this respect. With the same experimental setup, we assessed general autophagy using GFP-Atg8 and mitochondrial autophagy (mitophagy) using OM45-GFP. Autophagy in general, and mitophagy in particular, are induced rapidly upon introduction into sporulation media, even when cells are arrested prior to meiotic entry (figure S5E-G). Together these results indicate that cells perform autophagy throughout meiosis but prevent ERphagy until a later developmental stage. Because ERphagy has previously only been studied in the context of prolonged starvation or exposure to harsh chemical stress, it is intriguing to see its induction in a developmental context in which external stressors are absent.

### Meiotic ERphagy is mediated by selective autophagy receptors

Autophagy can occur either selectively or non-selectively. In selective autophagy, cargo-specific autophagy receptors recruit autophagosomes to their cargo via LC3-interacting region (LIR) motifs, ensuring precise target degradation (Anding and Baehrecke, 2017). Two budding yeast proteins, Atg39 and Atg40, have been identified as LIR motif-containing, ER-localized autophagy receptors (Mochida et al., 2015). During nitrogen starvation and rapamycin treatment, Atg39 mediates the autophagic degradation of the perinuclear ER and some nuclear material, whereas Atg40 promotes autophagy of the cortical ER. The developmental specificity of ERphagy induction suggests that cells exert some degree of selectivity in defining meiotic autophagy targets. To further determine if ERphagy in meiosis takes place selectively, we examined cells lacking either or both ERphagy receptors. Cells lacking Atg40 progressed normally through meiosis but autophagy of the cortical ER marker Rtn1-GFP was almost completely blocked (figure 6H-J). Atg39 was dispensable for degradation of cortical ER markers but important for autophagy of Sec63, a member of the translocon complex that localizes throughout the cortical and perinuclear ER (figure S5G-H). Thus, cells undergoing meiosis selectively target the ER for degradation via Atg39- and Atg40-mediated autophagy.

We also noted that ERphagy affects the overall abundance of target proteins over time, as Rtn1-GFP accumulates to much higher levels at late timepoints in autophagy-deficient *atg39Δ atg40Δ* mutants compared to WT (figure 6K). These results suggest that the remodeling of the ER proteome in late meiosis seen in our mass spectrometry data is driven, at least in part, by selective ERphagy.

### Atg40 expression is a developmental cue that triggers cortical ERphagy

ERphagy occurs primarily in late meiosis, downstream of the transcription factor Ndt80, but the precise developmental cues promoting ERphagy are still unclear (figure 6E-G). We examined Atg40 abundance during meiosis to determine if autophagy receptor expression itself might be the trigger for ER degradation. Atg40 was lowly expressed in vegetative cells and in early meiosis but was strongly induced in late meiosis, peaking in expression at around 6 hours before gradually declining (figure 6G, S5I). This spike in Atg40 levels immediately preceded the autophagic degradation of Rtn1, indicating that developmentally regulated autophagy receptor expression precisely coincides with the induction of autophagy.

Do other cues feed into ERphagy induction, or is autophagy receptor expression the principal regulatory cue? We noted that cells arrested prior to meiotic entry (*ime1Δ*) or in prophase I (*ndt80Δ*) show low Atg40 expression and exhibit very little ERphagy (figure 6E). To determine if providing cells with Atg40 in this context would be sufficient to induce ERphagy, we constructed a conditional allele of Atg40 using the copper-inducible *CUP1* promoter (*pCUP1-ATG40*). If Atg40 production is the limiting regulatory step for meiotic ERphagy, arrested cells should show Rtn1 degradation upon Atg40 induction. If additional developmental cues are required, Atg40 expression should be insufficient to trigger Rtn1 degradation. Consistent with the former model, copper-induced Atg40 expression resulted in robust autophagic degradation of Rtn1 during premeiotic arrest (figure 6L). In contrast, Atg40 overexpression in mitotic cells grown in rich media did not result in enhanced Rtn1 degradation. These results indicate that cells in sporulation media are primed for autophagy, and that the developmentally regulated expression of a single autophagy receptor is necessary and sufficient to trigger cargo degradation in this context.

### ER collapse is required for ERphagy

We noted that mutants with increased cortical ER retention in meiosis, namely *lnp1Δ* and *rtn1Δ rtn2Δ yop1Δ* cells, are also reported to be deficient in starvation-induced ERphagy (Chen et al., 2018). The converse is not true, as autophagy-deficient (*atg40Δ*) mutants display normal ER collapse (figure S6, video 28). We confirmed that *lnp1Δ* and *rtn1Δ rtn2Δ yop1Δ* mutant cells showed reduced ERphagy in the context of meiosis, which led us to investigate whether ER collapse is important for autophagic degradation in meiosis (figure 7A-B). One prediction of this model is that cortically retained ER is not subject to autophagy. Indeed, we saw very little autophagic degradation of the cortically retained tethers GFP-Ist2 and Tcb3-GFP relative to GFP-Scs2, an ER-PM tether that is not cortically retained, or Rtn1-GFP, a non-tether control (figure 7C).

**Figure 7.**
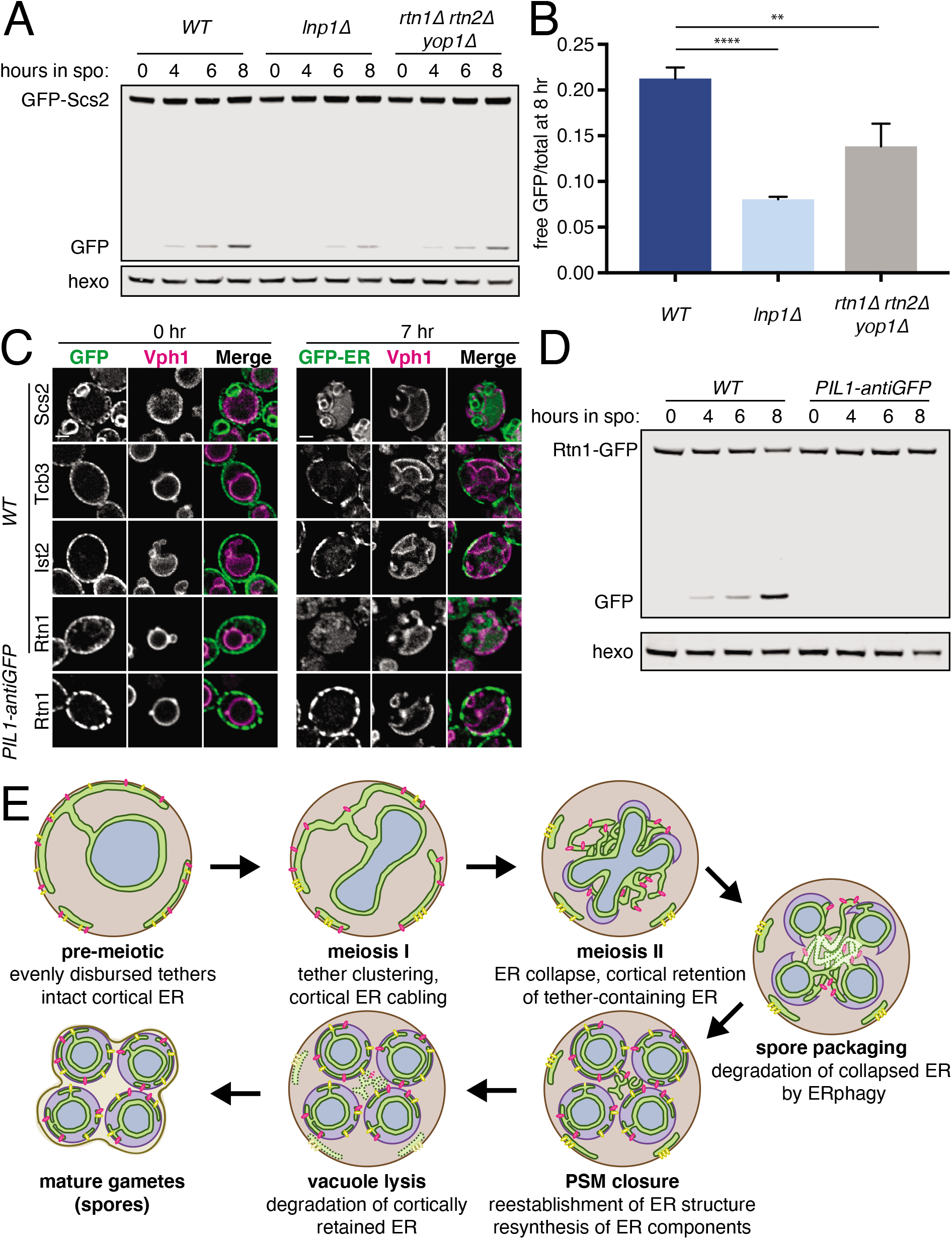
ER collapse is required for ERphagy. (A) Western blot with samples taken from cells of the indicated genotypes expressing GFP-Scs2 and Atg40-3V5 at the indicated times after transfer to sporulation media and probed for GFP and hexokinase. (B) Quantification of free GFP as a proportion of total GFP signal at 8 hours after transfer to sporulation media, using three replicates of the experiment from (A). (C) Microscopy images of cells expressing Vph1-mCherry and the indicated GFP-tagged ER protein and either an untagged (*WT*) or anti-GFP nanobody-tagged allele of Pil1 (*PIL1-antiGFP*). Images were taken at 0 and 7 hr following transfer to sporulation media. Scale bar = 2 μm. (D) Western blot with samples taken from cells expressing Rtn1-GFP and either WT Pil1 or Pil1-antiGFP. Samples were taken at the indicated times following transfer to sporulation media and probed for GFP and hexokinase. (E) Schematic showing key steps in meiotic ER remodeling.

If meiotic ERphagy depends on ER collapse, engineered targeting of a protein that is normally abundant in collapsed ER to the retained cortical ER compartment should no longer be subject to ERphagy. Indeed, ectopically targeting Rtn1-GFP to the cortically retained ER compartment using a Pil1-nanobody abolished autophagic degradation of Rtn1-GFP, supporting a model in which collapsed ER is robustly targeted by autophagy while cortically retained ER is not (figure 7C-D). Thus, the dramatic morphological changes in the ER during meiosis are integrally linked to its regulated degradation.

## Discussion

Here, we investigated the extensive ER remodeling that is a programmed into the natural developmental process of meiosis and gamete formation in budding yeast (figure 7E). This remodeling occurs in a stepwise manner, beginning early in meiosis with cortical ER cabling and ER-PM tether clustering. Next, coincident with anaphase II, the cortical ER undergoes reticulon- and Lnp1-dependent fragmentation and collapses from the plasma membrane, while a subset of ER fragments containing ER-PM tethers remain at the cell cortex. Collapsed ER is then subject to Atg40-dependent selective autophagy, whereas cortically retained ER is degraded by following spore closure and vacuole lysis. Together these findings reveal that developmentally regulated ER compartmentalization, membrane tethering, and two parallel pathways for ER degradation combine to mediate ER inheritance by gametes while selectively eliminating a subset of ER.

An especially unexpected finding presented here is the meiotically regulated transition of cortical ER from one continuous membrane system to separate fragmented topologies, with the fate of each fragment determined by whether or not it contains ER-PM tethers. Tether-free cortical ER is subject to ER collapse, whereas tether-containing fragments remain at the plasma membrane and are excluded from gametes. Consistent with this model, artificially tethering the abundant cortical ER protein Rtn1 to the PM increases cortical ER retention but does not prevent ER collapse. Although programmed ER fragmentation has not, to our knowledge, been described during differentiation, it elegantly achieves two distinct outcomes at once, allowing bulk cortical ER inheritance while partitioning specific ER fragments away from gametes.

A major unanswered question is how reticulon-mediated membrane fragmentation is regulated to achieve precisely timed ER detachment. A large body of evidence suggests that increased reticulon concentration drives membrane fission. In some cases, fragmentation occurs globally upon reticulon overexpression (Espadas et. al., 2019; Wang et. al., 2016), whereas in others fragmentation relies a high *local* reticulon concentration achieved through multivalent interactions or homo-oligomerization (Mochida et. al., 2020; Bhaskara et. al., 2019; Jiang et. al., 2020; Grumati et. al., 2017). In at least one case, phosphorylation of a reticulon-like protein promotes oligomerization-dependent fragmentation (Jiang et. al., 2020). Several lines of reasoning lead us to favor a regulated oligomerization model rather than increased reticulon expression as the mechanism driving meiotic ER fragmentation. First, new protein synthesis on the scale required to drive fragmentation is slow and energetically costly relative to the regulatory mechanisms that might control oligomerization, such as phosphorylation. Second, global measurements of protein synthesis and abundances suggest that reticulon levels do not appreciably increase during the time leading up to ER collapse (Brar et. al., 2012; Cheng et. al., 2018). Third, fragmentation purely as a result of reticulon abundance has only been observed in *in vitro* or overexpression systems in which reticulons exceed physiological levels. In contrast, regulated reticulon oligomerization is a logical and concerted mechanism deployed by cells in response to specific challenges (Jiang et. al., 2019). Our finding that yeast reticulons are involved in meiotic ER collapse and inheritance further highlights that these proteins are more than inert structural proteins, and instead can serve critical roles in dynamic ER remodeling. It will be important to determine the developmental cues regulating reticulon-dependent membrane scission in meiosis, and whether coordinated reticulon oligomerization drives this phenomenon.

Morphological homeostasis of the tubular ER network requires a balance between membrane fusion and fission. We propose that this balance shifts toward fission in meiotic cells, resulting in fragmented ER. Consistent with a need for reduced tubule fusion in this process, we found that Lnp1, which has been suggested as a negative regulator of Sey1-mediated tubule fusion (Chen et. al., 2012), is important for ER collapse in meiosis. Cells lacking Lnp1 formed massive ER foci that remained attached to the cell cortex throughout meiosis. How these structures form and their precise composition are still unclear, though the decrease in focus size and abundance in *lnp1Δ sey1Δ* double mutant cells suggests that they are the result of excessive membrane fusion in the absence of Lnp1. These structures are not present in *lnp1Δ* cells undergoing exponential mitotic growth, though we did observe them in saturated *lnp1Δ* cultures (data not shown), suggesting an unexpected role for Lnp1-dependent ER remodeling during nutrient adaptation. Further elucidation of the function of Lnp1 in these conditions may provide crucial insight into the role of this conserved yet poorly understood family of proteins.

We found that four ER-PM tethers, namely Ist2 and the three tricalbins, mark cortically retained ER and are important for the physical exclusion of these ER fragments from gametes. Deletion of these tethers reduces the amount of cortically retained ER at anaphase II but does not completely prevent ER retention, suggesting that additional tethers and/or alternative mechanisms ensure cortical ER tethering in this context. While Scs2 and Scs22 are normally released from the plasma membrane during meiosis, they may contribute to cortical ER retention in the absence of the other four tethers. However, deletion of all six tethers eliminates almost all cortical ER even in non-meiotic cells, complicating our ability to assess meiotic cortical ER retention in this genotype. In addition to these six tethers, other proteins have been observed to localize to ER-PM contact sites, and some have been proposed as active ER-PM tethering proteins (Petkovic et al., 2014; Topolska et al., 2020). Study of these additional factors will be important to further interrogate the molecular basis of cortical ER retention during meiosis and its role in ensuring gamete health.

What is the purpose of excluding certain parts of the ER from inheritance by gametes? One possibility is that this process serves as a form of ER quality control, selectively preventing the inheritance of damaged or toxic ER or ER-associated material. Targeting deleterious ER contents to the cortically retained compartment would be an efficient means of ensuring their physical exclusion from gametes. An important challenge with such a system would be achieving specificity in what is targeted to cortically retained ER. Thus far, the only proteins that we identified to preferentially localize to the cortically retained compartments are ER-PM tethers themselves. It will be important to determine in future studies whether other ER proteins or cellular components selectively localize to this compartment, and the role that ER-PM tethers play in this process.

In addition to the exclusion of cortically retained ER from gametes, we identified programmed selective autophagy as a parallel and mutually exclusive means of eliminating ER subdomains during meiosis. Although ERphagy has primarily been studied in the context of nutrient limitation or ER stress in the presence of harsh chemical treatment, our identification of its natural role in meiosis provides an opportunity to study its endogenous regulation in a developmental context. We identified the timed expression of the ER-specific autophagy receptor Atg40 as a key developmental cue regulating ERphagy in meiosis. Atg40 expression depends on the meiotic transcription factor Ndt80, but whether Ndt80 promotes Atg40 expression directly remains to be determined. Previous work has shown that components of the histone deacetylase complex Rpd3L repress *ATG40* transcription in nutrient-replete conditions (Cui et al., 2019), suggesting that inhibition of this complex may allow meiotic Atg40 expression. Our findings warrant a more detailed study of how its expression and activity are regulated, and the role of ERphagy in the broader developmental context of meiosis.

As with cortical ER retention, it is appealing to hypothesize that ERphagy serves as a meiotic quality control mechanism. The autophagy receptors Atg39 and Atg40 are selective for perinuclear and cortical ER, respectively, during starvation-induced and meiotic ERphagy. However, whether there is additional specificity to what ER content is degraded in any context is yet to be determined. ERphagy receptors in mammalian cells have been shown to preferentially degrade subsets of the ER proteome, including misfolded proinsulin and procollagen aggregates, but the molecular basis for this specificity is largely unknown (An et al., 2019; Cunningham et al., 2019; Forrester et al., 2019). In the future, understanding how cells precisely target cargo for degradation by ERphagy to remodel the ER and mitigate ER stress will be crucial for defining the role of ERphagy meiosis. Moreover, owing to the conservation of ERphagy factors in mammals, we propose that the natural role of ERphagy in meiotic development suggests a more widespread role for this newly uncovered process in mammalian cellular development programs.

The most widely studied substrates for ER quality control are misfolded proteins, which are induced by genetic or chemical disruption of ER protein folding capacity, or through exogenous expression of model aggregate-prone proteins. During mitosis in budding yeast, misfolded ER proteins are retained in mother cells, promoting daughter cell rejuvenation at the expense of reduced mother cell lifespan (Clay et al., 2014; Piña and Niwa, 2015). Intriguingly, while the vast majority of ER components enter into the daughter cell during mitotic ER inheritance in unstressed conditions, all four cortically retained tethers are tightly restricted to the mother cell (Okada et al., 2017; Sugiyama and Tanaka, 2019; Takizawa et al., 2000). Moreover, asymmetric inheritance of both misfolded proteins and ER-PM tethers relies on an ER membrane diffusion barrier established by septin ring components at the bud neck (Clay et. al., 2014; Sugiyama and Tanaka, 2019), suggesting a shared mechanism to control the selective inheritance of ER protein aggregates and ER-PM tethers, and raising the possibility that ER-PM tethers themselves may mark or actively participate in the age-dependent accumulation of ER stress. A conserved but understudied feature of gametogensis is the elimination of age-induced damage to produce healthy, youthful gametes (Goodman et. al., 2020). A variety of abnormal structures, including cytosolic protein aggregates, extrachromosomal rDNA circles, and expanded nucleoli, accumulate with age in budding yeast and are retained by mother cells during cell division (Shcheprova et al., 2008). During gamete formation, most of these structures are excluded from gamete cells and eliminated, likely contributing to gamete rejuvenation (King et al., 2019; Ünal et al., 2011). While naturally occurring markers of age-induced ER damage have not been described in yeast, our work motivates further study of relationship between aging, ER stress, and cortical ER inheritance during meiosis and mitosis.

## Methods

### Yeast strains, plasmids, and primers

All experimental strains are diploid *Saccharomyces cerevisiae* derivatives of the SK1 strain as detailed in supplemental table 1. The following alleles were derived from previous studies: *pGAL-NDT80* and *GAL4.ER* (Benjamin et al., 2003; Carlile and Amon, 2008), *pCLB2-CDC20* (Lee and Amon, 2003), *pCUP1-oSTIR* (Sawyer et al., 2019), *mKate-SPO20_51-91_* and *VPH1-mCherry* (King et. al., 2019), *HTB1-mCherry* (Matos et al., 2008), *ndt80Δ* (Xu et al., 1995), *GFP-ATG8* (Graef et al., 2013). The *atg8::LEU2* allele was a gift from Hilla Weidberg and Angelika Amon.

Deletion and C-terminal tagging of genes at their endogenous loci were performed using an established PCR-based technique (Janke et al., 2004; Longtine et al., 1998) using the primers indicated in supplemental table 2 and plasmids indicated in supplemental table 3. *GFP-SCS2* and *GFP-IST2* were created using the Cas9-based method described in Sawyer et. al. (2019) using guide RNAs detailed in supplemental table 2. The repair template containing GFP and a linker sequence was amplified from pÜB1548. GFP-Scs22 expressed from its endogenous locus was not detectable by microscopy in our strain background (not shown), so we generated an allele that is highly expressed in meiosis (*pATG8-GFP-SCS22). SCS22* was amplified from SK1 genomic DNA and cloned into pÜB 1548 by Gibson assembly (Gibson et al., 2009), replacing the *ATG8* ORF. The *SCS22* intron was removed following the Q5 Site-Directed Mutagenesis protocol (New England Biolabs) to generate pÜB1889. This plasmid was digested with PstI and transformed into WT SK1. To generate *pCUP1-ATG40-3V5, ATG40-3V5* was amplified from yeast harboring that allele and cloned together with the *CUP1* promoter into the pÜB217 plasmid by Gibson assembly. The resulting plasmid was amplified with the primers indicated in Table 2 and transformed into a strain harboring *atg40::KanMX*, replacing the KanMX cassette to give *atg40::pCUP1-ATG40-3V5-HygB*. To construct a *GFP-HDEL* construct that is stably expressed throughout meiosis, the *GFP-HDEL* sequence published in Rossanese et al., 2001 (2001) was cloned into a *TRP1* integrating vector harboring the *ARO10* promoter, obtained from Leon Chan. The resulting *pARO10-GFP-HDEL-TRP1* construct was used to generate *pARO10-mCherry-HDEL* by Gibson assembly. Both constructs were transformed into yeast following digestion with PmeI.

### Media and growth conditions

Prior to the induction of meiosis, cells were grown at room temperature for 20-24 hr to a density of OD_600_ ≥ 10 in YEPD (1% yeast extract, 2% peptone, 2% glucose, 22.4 mg/L uracil, 80 mg/L tryptophan). Cultures were then diluted to OD_600_ = 0.25 in BYTA (1% yeast extract, 2% bacto tryptone, 1% potassium acetate, and 50 mM potassium phthalate) and grown for 16-18 (OD_600_ ≥ 4.5) hr at 30° C. Cells were then pelleted, washed with sterile MilliQ water and resuspended to OD_600_ = 1.9 in sporulation medium (SPO; 2% potassium acetate, 40 mg/L adenine, 40 mg/L uracil, 10 mg/L histidine, 10 mg/L leucine and 10 mg/L tryptophan adjusted to pH 7.0 and supplemented with 0.02% raffinose). Cultures were allowed to shake at 30° C for the duration of the experiment. For each stage, culture volume was 1/10^th^ of the flask volume to ensure proper aeration.

For experiments conducted during vegetative growth, cells were grown in YEPD for 16-18 hours at 30° C (OD_600_ ≥ 10). Cultures were then back-diluted to OD_600_ = 0.02-0.05. For imaging experiments, cells were examined at a density of OD_600_ = 0.6-0.8.

For experiments using the *pCUP1-ATG40-3V5 allele*, CuSO_4_ was added to a final concentration of 50 μM at the indicated times. For *pGAL-NDT80* experiments, β-estradiol or an equivalent volume of 100% EtOH was added to a final concentration of 1 μM β-estradiol. For Atg1-AID experiments, 50 μM CuSO_4_ was added to induce expression of *pCUP1-osTIR* followed immediately by 500 μM auxin (Sigma).

### Live-cell imaging

Live-cell imaging was performed exactly as described in King et. al. (2019), except fresh SPO was used in place of conditioned SPO. Specific imaging conditions are noted in supplemental table 4. All time-lapse experiments were performed using the CellAsic system (EMD Millipore) in Y04D or Y04E microfluidics plates, with the exception of the Latrunculin A experiments, for which we used glass-bottom 96-well plates (Corning).

### Image quantification

For time-lapse microscopy, anaphase I was defined as the first frame in which an elongated nucleus was observed (if applicable), or the first frame at which two distinct nuclear masses were visible. Anaphase II was defined as the first frame at which two elongated nuclei were observed following anaphase I. ER cabling was defined as the first frame at which ER cables were visible ER cables are bright, cortically localized ER structures that are thicker and more dynamic than pre-meiotic cortical ER. ER collapse was defined as an abrupt movement of cortical ER toward the center of cells. Prospore membrane nucleation was defined as the first frame at which mKate-Spo20_51-91_ signal was visible as distinct puncta in the center of cells rather than plasma membrane-localized. Prospore membrane closure was defined as the frame at which membrane structure transitioned from elongated to circular. Vacuole lysis was defined as the time at which signal became diffuse rather than membrane-localized. Degradation of GFP-Ist2 and Tcb3-GFP was defined as the frame at which their signal disappeared from the cell cortex.

Qualitative cortical ER classification was performed at anaphase II according to the guidelines outlined in figure 2D. Class III ER (“intermediate”) includes cells that had either a mixture of small fragments and large spans of cortical ER AND/OR fragments that were intermediate in size and therefore not categorized as small fragments or large spans.

For Gini index calculation, the cell periphery was traced for the centermost Z-slice using the program Fiji. Pixel intensity was calculated along the length of the trace, resulting in a finite number of measurements “n”. These measurements were then ordered from smallest to largest and given an integer ranking “i” based on this order (i.e. for each value 1 ≤ i ≤ n, where the smallest number in the dataset has i=1 and the highest number has i=n). Background was subtracted using average pixel intensity from a cell-free region of the image. The Gini index (G) was determined using the formula:

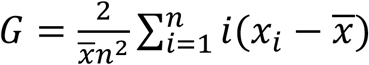

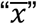 is the average of all measurements and xi is the intensity value of ranking “i” in the dataset. For each cell (n=10), Gini values were calculated for at least seven timepoints prior to ER collapse and six timepoints following ER collapse.

For analysis of foci in *lnp1Δ* cells, foci were counted manually for at least 100 cells per genotype. Focus size was measured using Fiji. Briefly, images were z-projected using maximum intensity projection, converted to 8-bit, and threshold was adjusted so that foci were clearly visible. Foci were detected automatically using the “analyze particles function, resulting in measurements for at least 134 foci across over 100 cells per genotype.

For measurements of Rtn1 and Htb1 levels in heterozygously tagged cells, images were maximum projected over the full imaging volume in Fiji. Tracing was performed for the whole cell (anaphase I and anaphase timepoints) or for individual spores, and average pixel intensity for the traced area was calculated for both channels. Measurements for the cells shown in figure 5C-F were obtained from the first frame at which individual spores were easily distinguishable until spores became tightly packed and therefore had significantly overlapping signal (at least 480 min). Bright spore and dim spore images in figure 5G were taken from the last timepoint at which spores did not significantly overlap.

### Meiotic staging

Meiotic staging was performed scoring DAPI and tubulin morphology by fluorescent microscopy as described in Sawyer et. al. (2019). Samples were fixed in 3.7% formaldehyde for 12-24 hr at 4° C. Cells were then washed with 100 mM potassium phosphate pH6.4, once with sorbitol citrate (100 mM potassium phosphate pH7.5, 1.2 M sorbitol), and digested in 200 μL sorbitol citrate, 20 μL glusulase (Perkin-Elmer) and 6 μL of zymolase (10 mg/mL, MP Biomedicals) for 3 hours at 30° C while rotating. Samples were pelleted at 900 rcf for 2 min, washed with 100 μL sorbitol citrate, pelleted again and resuspended in 50 μL sorbitol citrate. Samples were then mounted on slides prepared with poly-L-lysine, submerged in 100% methanol at −20° C for 3 min, transferred to 100% acetone at −20° C for 10 sec, then allowed to air dry. Samples were then incubated at RT for 1 hr in primary anti-tubulin antibody (Bio-Rad, 1:200) in PBS-BSA (5 mM potassium phosphate, 15 mM NaCl, 1% BSA, 0.1% sodium azide). Samples were then washed 3x in PBS-BSA and incubated with preabsorbed FITC-conjugated secondary antibody (Jackson ImmunoResearch Labs, 6:200) for 1 hr at RT. Samples were washed 3x with PBS-BSA and mounted with VectaShield Antifade Mounting Medium with DAPI (Vector Labs).

### Western blotting

Protein samples were extracted by trichloroacetic acid (TCA) precipitation as described previously (Chen et al., 2017), with some modifications. For meiotic samples, 1.8 mL culture was mixed with 200 μL 50% TCA (5% final concentration) and incubated at 4° C for 12-24 hr. For vegetative mitotic samples, 3.42 ODU culture was spun down for 2 min at 3000 rcf, washed in sterile MilliQ water, resuspended in 5% TCA and incubated at 4° C for 12-24 hr. All samples were precipitated for 5 min at 20,000 rcf and washed in 1 mL acetone. The acetone was aspirated and samples were allowed to dry for at least 20 minutes. Pellets were resuspended by bead beating for 5 min in 100 μL TE supplemented with 3 mM and 1x protease inhibitors (Roche) with 100 μL acid-washed glass beads. 50 μL 3x SDS loading buffer was added and samples were incubated at 95° C for 5 minutes and spun down for 5 min at 20,000 rcf. 4 μL were loaded onto a Bis-Tris acrylamide gel, separated at 150 V for 50 minutes and transferred to a nitrocellulose membrane using the TransBlot Turbo system (BioRad). Blots were blocked and probed overnight at 4° C with one or more of the following antibodies: anti-hexokinase (US Biological, 1:15,000), anti-GFP JL8 (Clontech, 1:2000), anti-V5 (Invitrogen, 1:2000). Blots were washed in PBST and incubated for 2 hr in IRDye secondary antibodies (LI-COR, 1:20,000). Blots were imaged and quantified using the Odyssey system (LI-COR).

## Acknowledgements

We thank Elçin Ünal, James Olzmann, Roberto Zoncu, Ina Hollerer and Tina Sing for their comments on this manuscript. We also thank Max Ferrin, Yidi Sun, and David Drubin for generously sharing reagents and technical advice; and Grant King and Elçin Ünal for sharing data and technical advice. This work was supported by Rose Hills Foundation and National Institutes of Health [1R35GM134886] funding to G.A.B. G.M.O was funded in part by NIH training grants to the MCB department (T32 GM 0007232 and T32 HG 00047).

---

**Figure S1.**
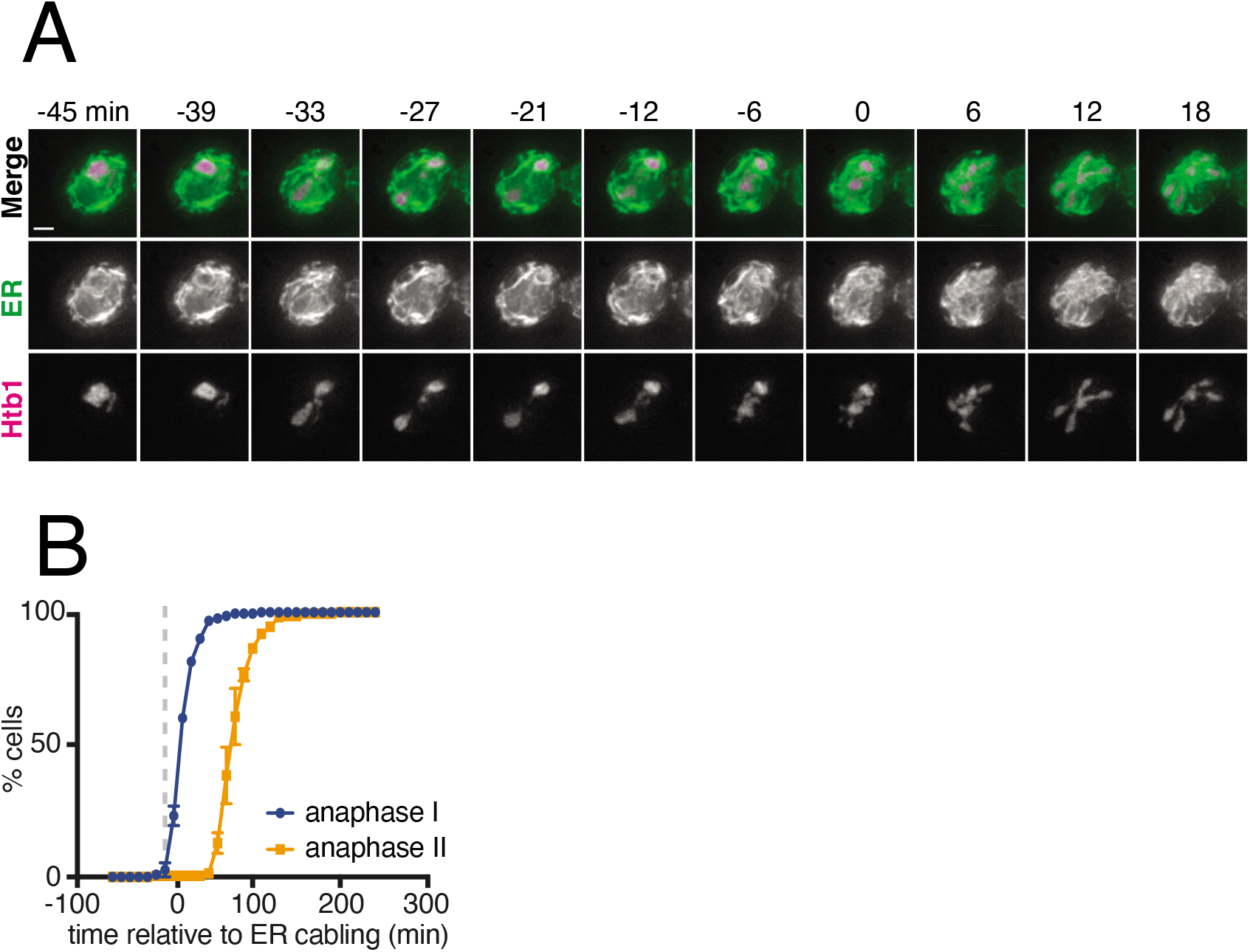
The ER undergoes developmentally regulated structural remodeling during meiosis. (A) Time-lapse microscopy of cells expressing GFP-HDEL (ER) and Htb1-mCherry imaged every 3 minutes in meiosis. Minute 0 is defined as the time of ER collapse. (B) Quantification of the time of anaphase I and anaphase II relative to ER cabling.

**Figure S2.**
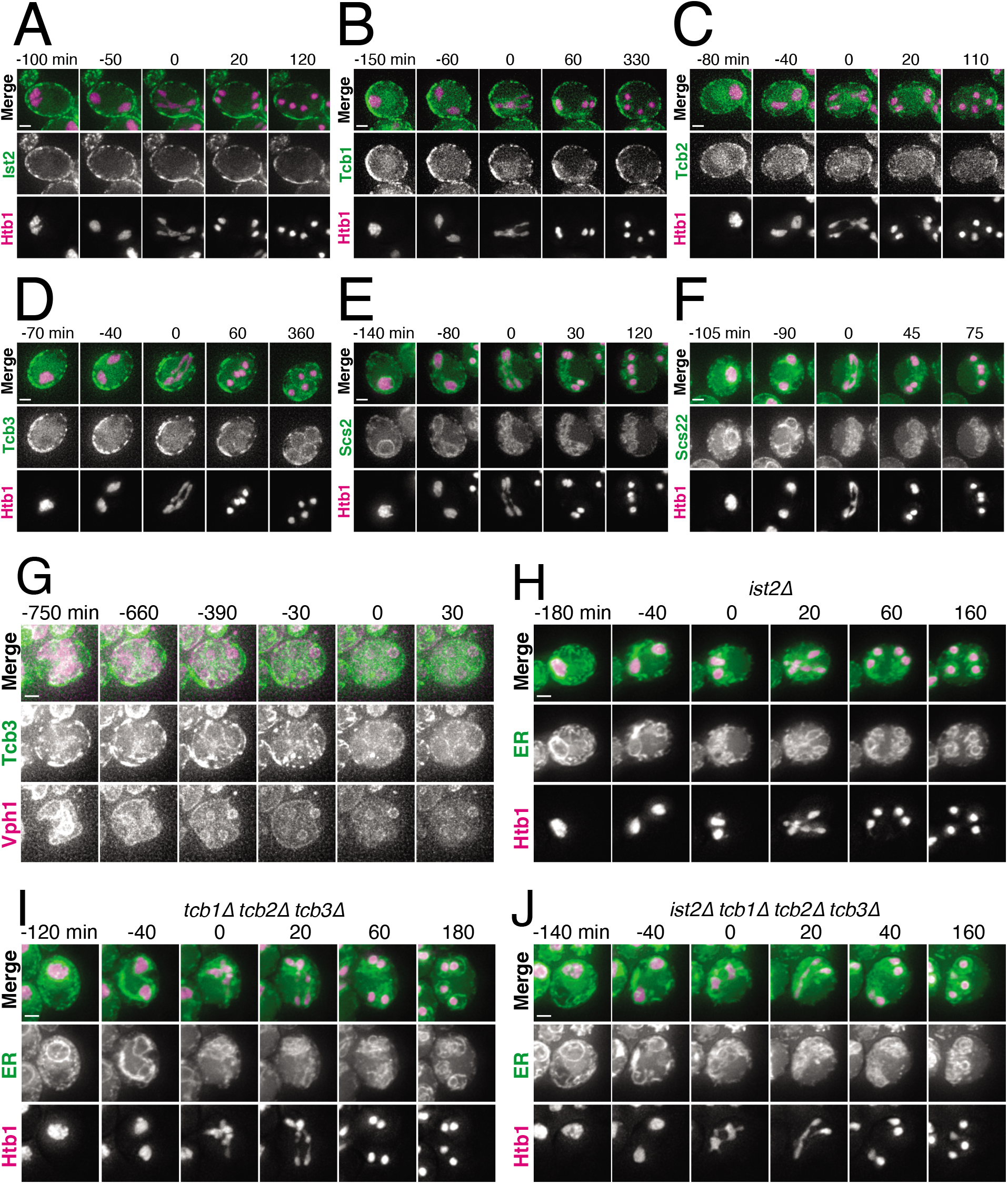
A subset of ER-PM tethering proteins marks cortically retained ER islands. (A-F) Time-lapse microscopy of cells expressing Htb1-mCherry and the indicated GFP-tagged ER-PM tether protein, imaged every 10 minutes during meiosis. Minute 0 is defined as the onset of anaphase II. (G) Time-lapse microscopy of cells expressing Tcb3-GFP and Vph1-mCherry imaged every 30 minutes during meiosis. Minute 0 is defined as the time of vacuole lysis. (H-I) Time-lapse microscopy of cells of the indicated genotypes expressing GFP-HDEL (ER) and Htb1-mCherry imaged every 10 minutes during meiosis. Minute 0 is defined as the time of ER collapse. Scale bar = 2 μm for all panels.

**Figure S3.**
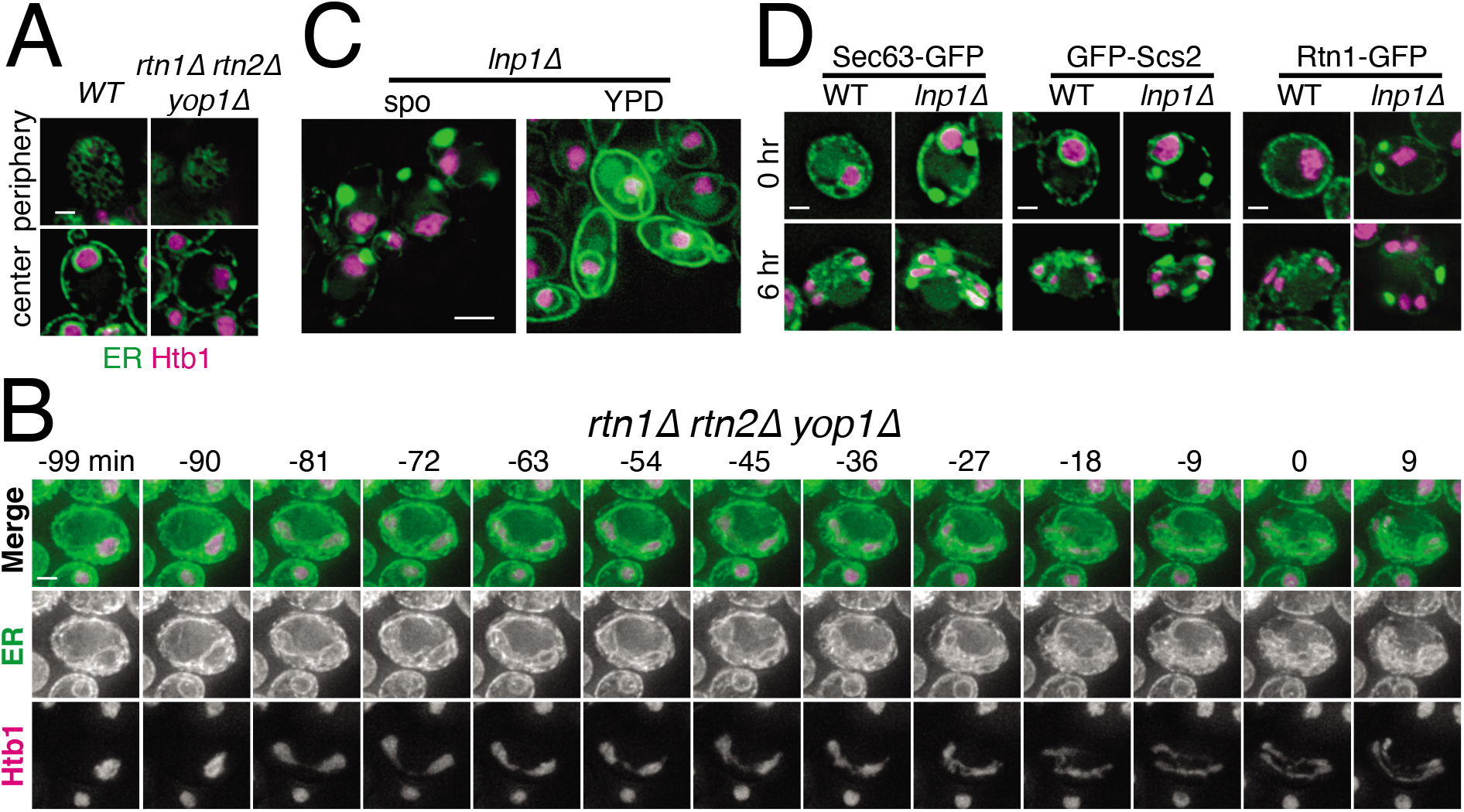
Reticulons and Lnp1 regulate meiotic ER remodeling. (A) WT and *rtn1Δ rtn2Δ yop1Δ* cells expressing GFP-HDEL (ER) and Htb1-mCherry imaged at the cell periphery and cell center immediately after transfer to sporulation media. (B) Time1apse microscopy of *rtn1Δ rtn2Δ yop1Δ* cells expressing GFP-HDEL (ER) and Htb1-mCherry imaged every 3 minutes in meiosis. Minute 0 is defined as the time of ER collapse. (C) *lnp1Δ* cells expressing GFP-HDEL (ER) and Htb1-mCherry imaged immediately following transfer to sporulation media, or during exponential growth in YPD. (E) WT and *lnp1Δ* cells expressing Htb1-mCherry and the indicated GFP-tagged protein imaged at 0 or 6 hr after transfer to sporulation media. Scale bar = 2 μm for all panels.

**Figure S4.**
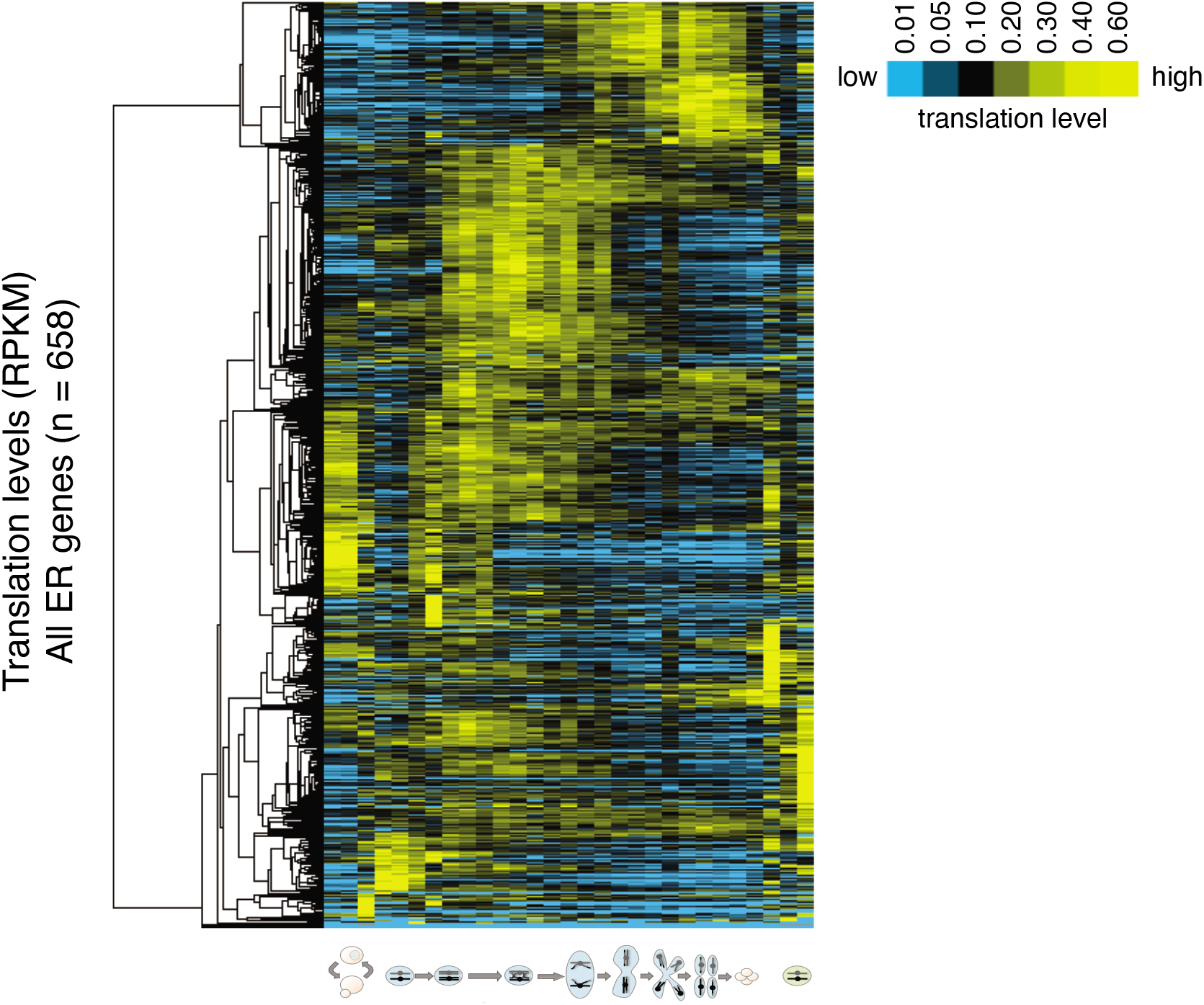
The ER undergoes turnover and resynthesis during meiosis. Hierarchical clustering of translation levels for all genes annotated for ER localization using on ribosome profiling data published in Brar, *et. al*. (2012). Footprint RPKM are normalized within each gene to facilitate comparisons of expression patterns between genes independent of overall abundance.

**Figure S5.**
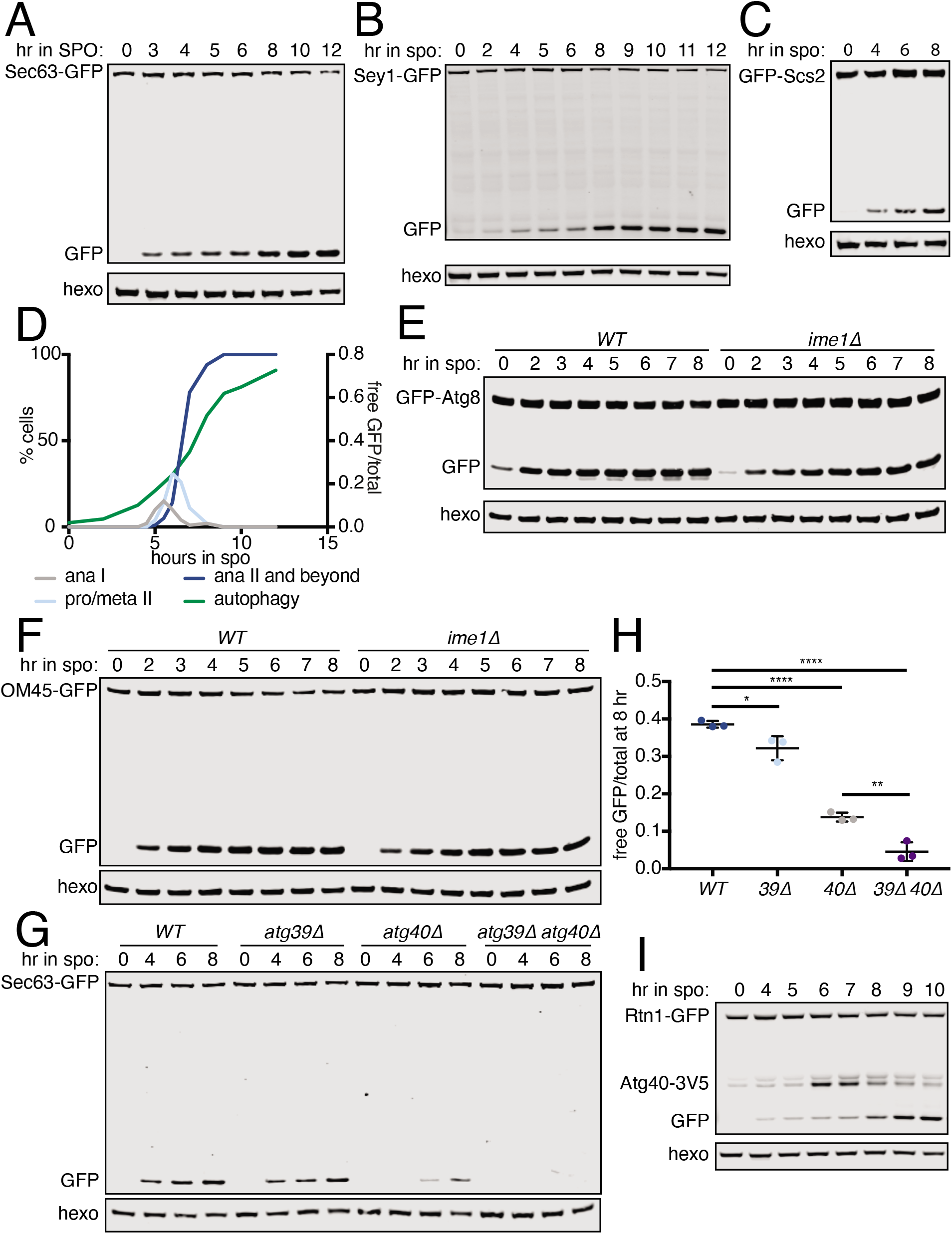
The ER is degraded by selective autophagy during meiosis. (A-C) Western blot using samples from WT cells expressing the indicated GFP-tagged protein taken at the indicated times in meiosis. Blots were probed for GFP and hexokinase. (D) Quantification of meiotic staging and autophagy using samples taken in parallel to those in figure 6A. Left axis shows the % of cells at the indicated stage in meiosis and right axis shows the free GFP signal as a proportion of the total (Rtn1-GFP + free GFP). (E) Western blot using samples from WT and *ime1Δ* cells expressing GFP-Atg8 taken at the indicated times during meiosis. Blots were probed for GFP and hexokinase. (F) As in (E) but with cells expressing OM45-GFP instead of GFP-Atg8. (G) Western blot using samples from cells of the indicated genotypes expressing Sec63-GFP taken at the indicated times in meiosis. Blots were probed for GFP and hexokinase. (H) Quantification of free GFP as a proportion of the total GFP signal at 8 hr from three replicates of the experiment in (G). (I) Western blot using samples from WT cells expressing Rtn1-GFP and Atg40-3V5 taken at the indicated times in meiosis. Blot was probed for GFP, V5 and hexokinase.

**Figure S6.**
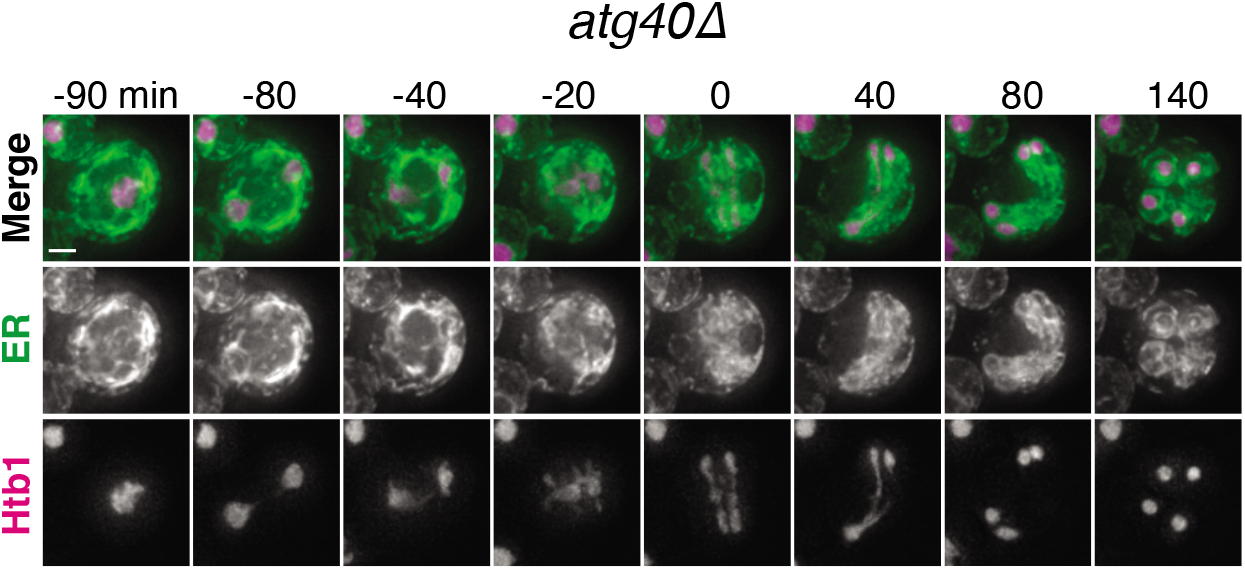
ER collapse is required for ERphagy. Time-lapse microscopy of *atg40Δ* cells expressing GFP-HDEL (ER) and Htb1-mCherry imaged every 10 minutes during meiosis. 0 min is defined as the time of ER collapse. Scale bar = 2 μm.

**Table S1.**
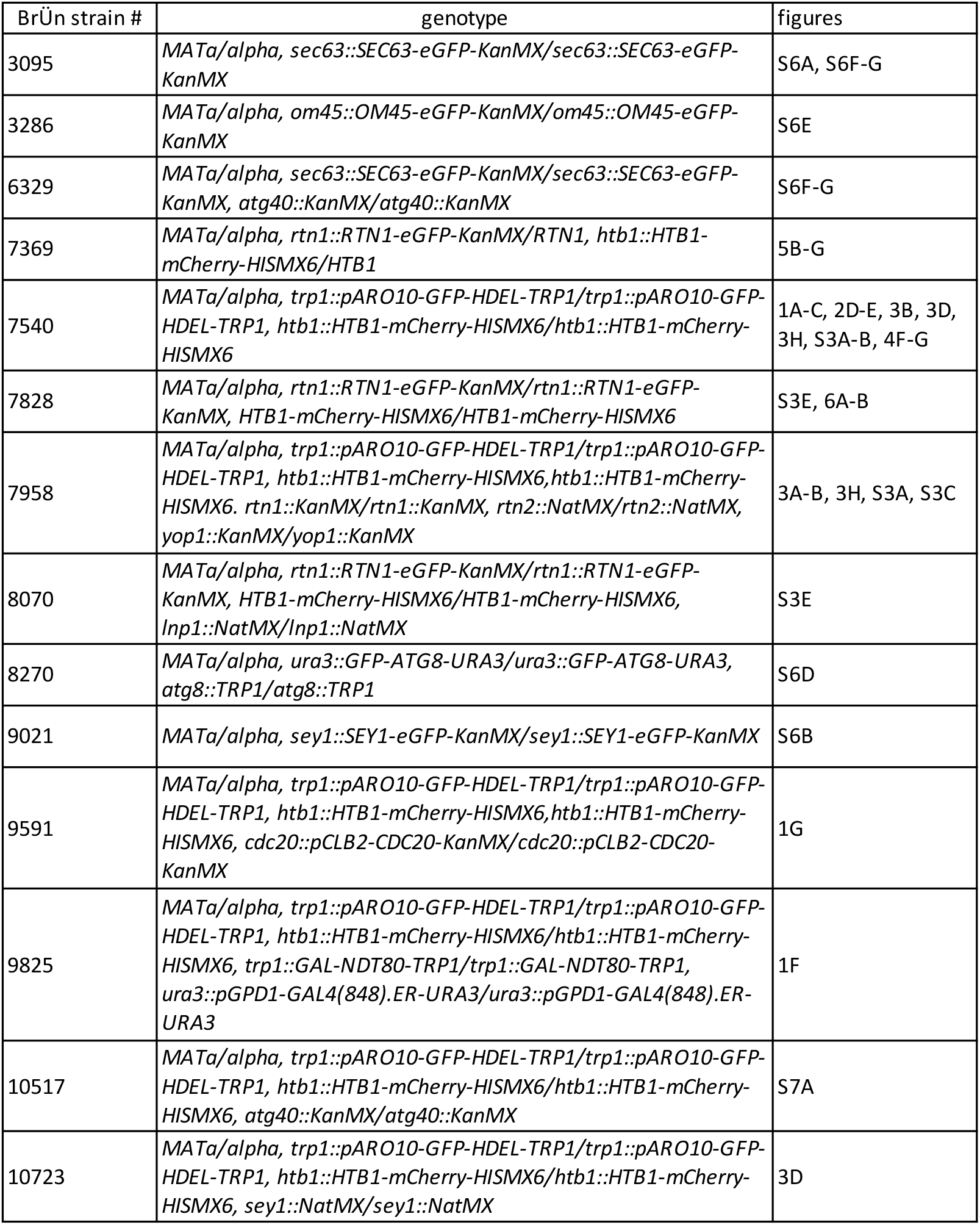
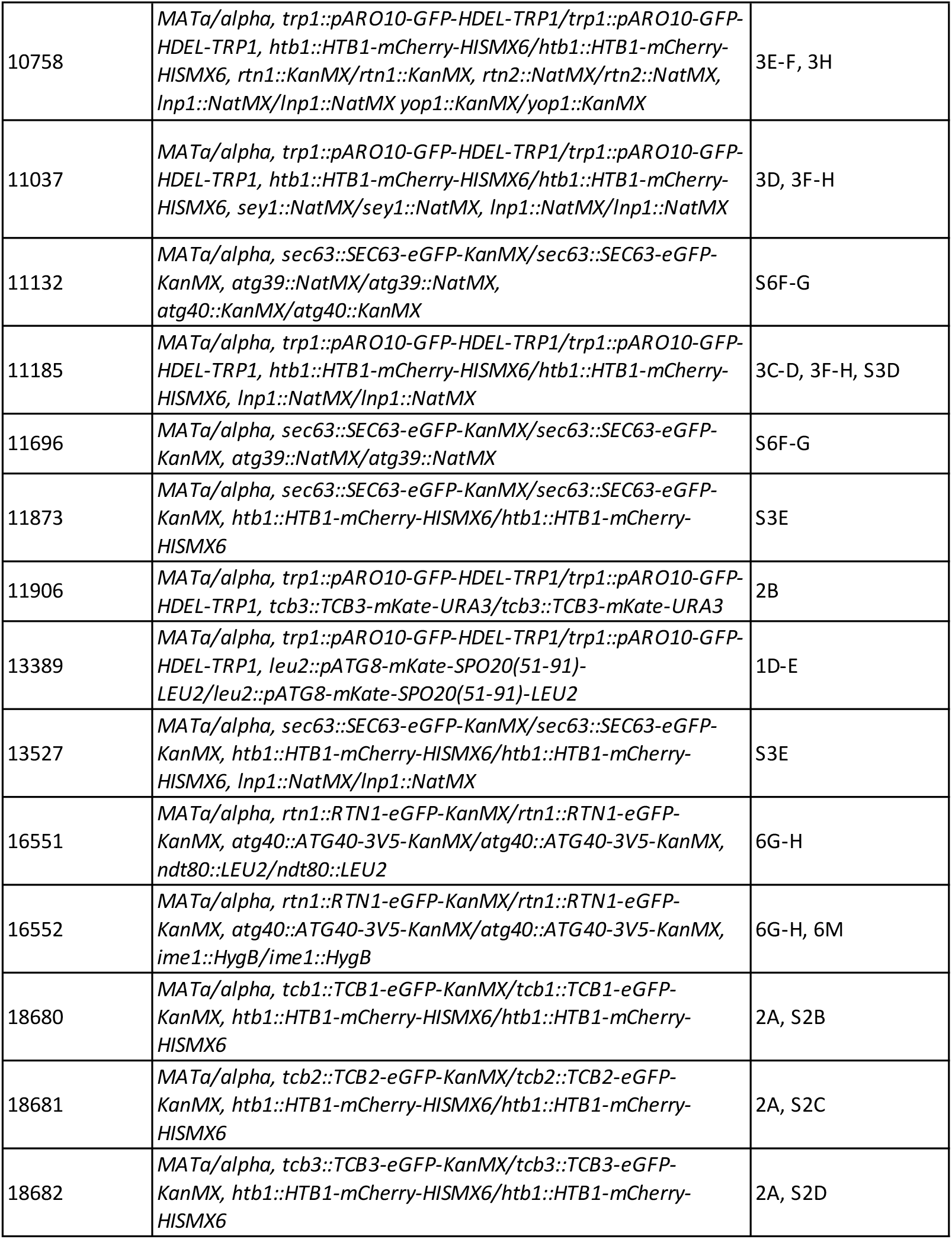
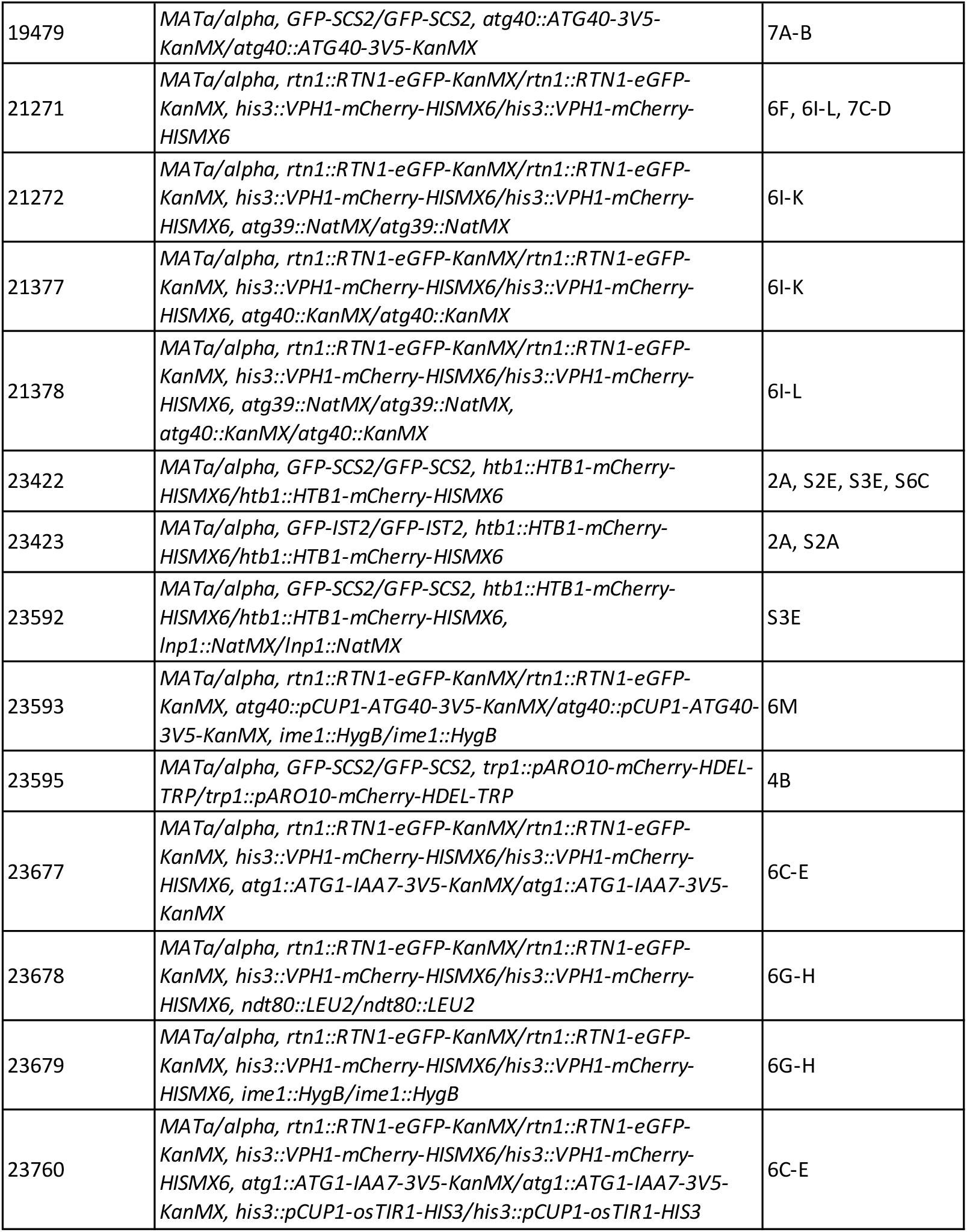
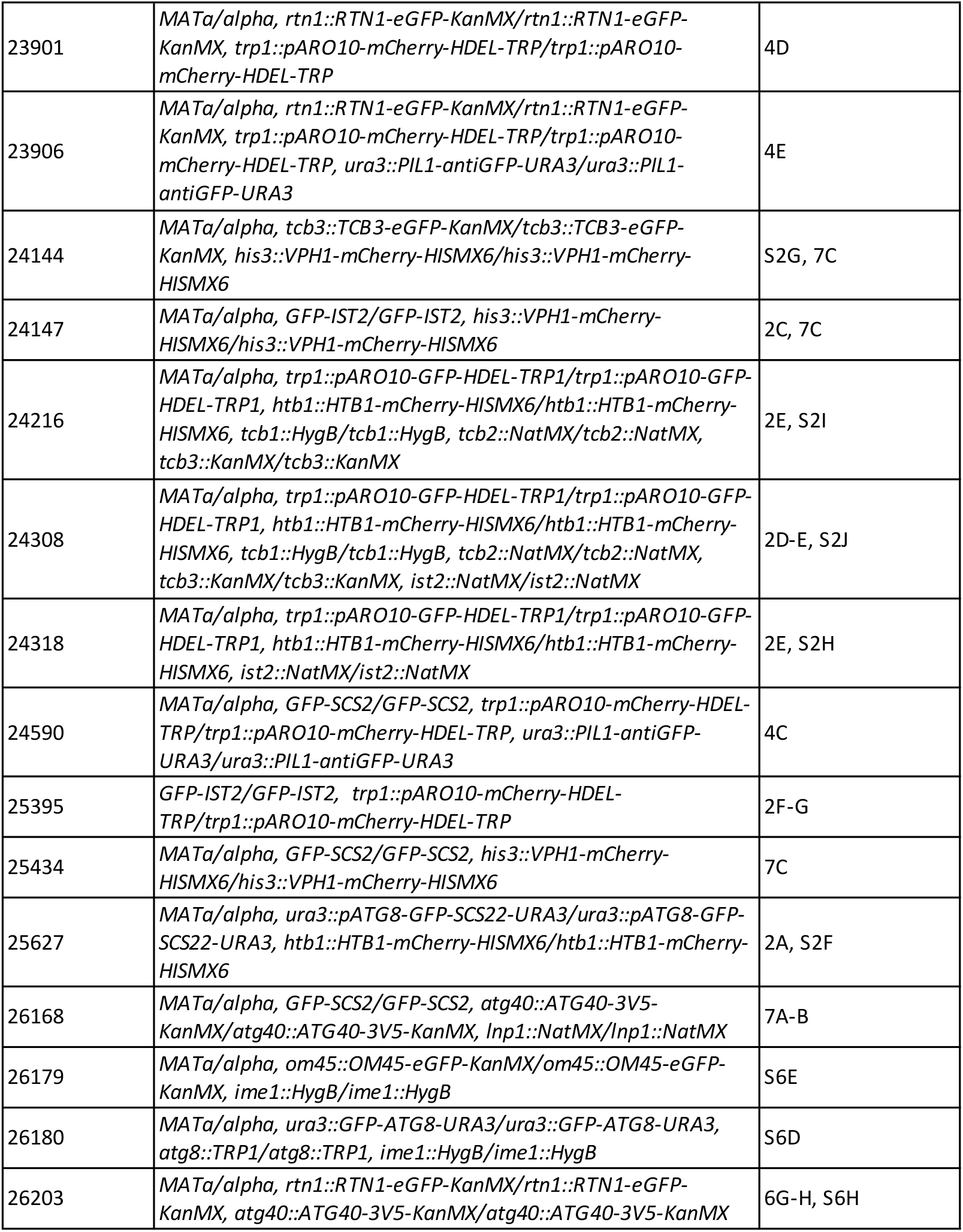
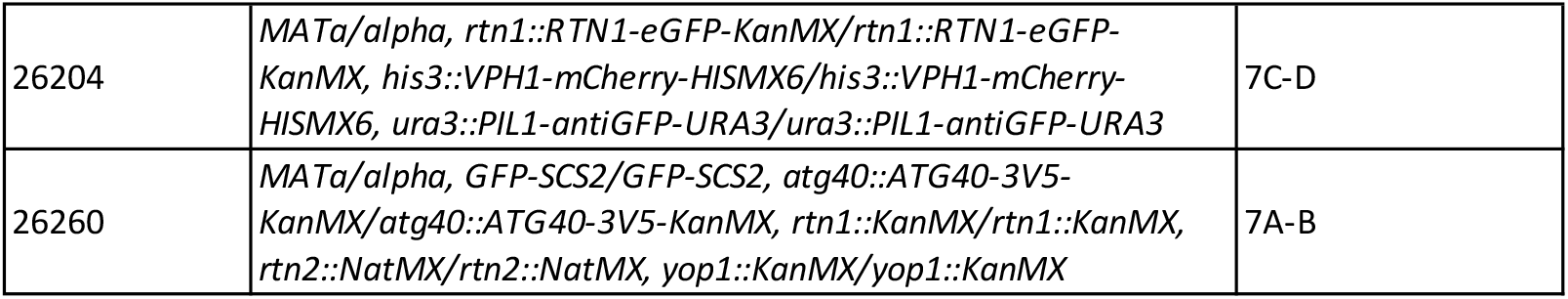
Strains used in this study.

**Table S2.**
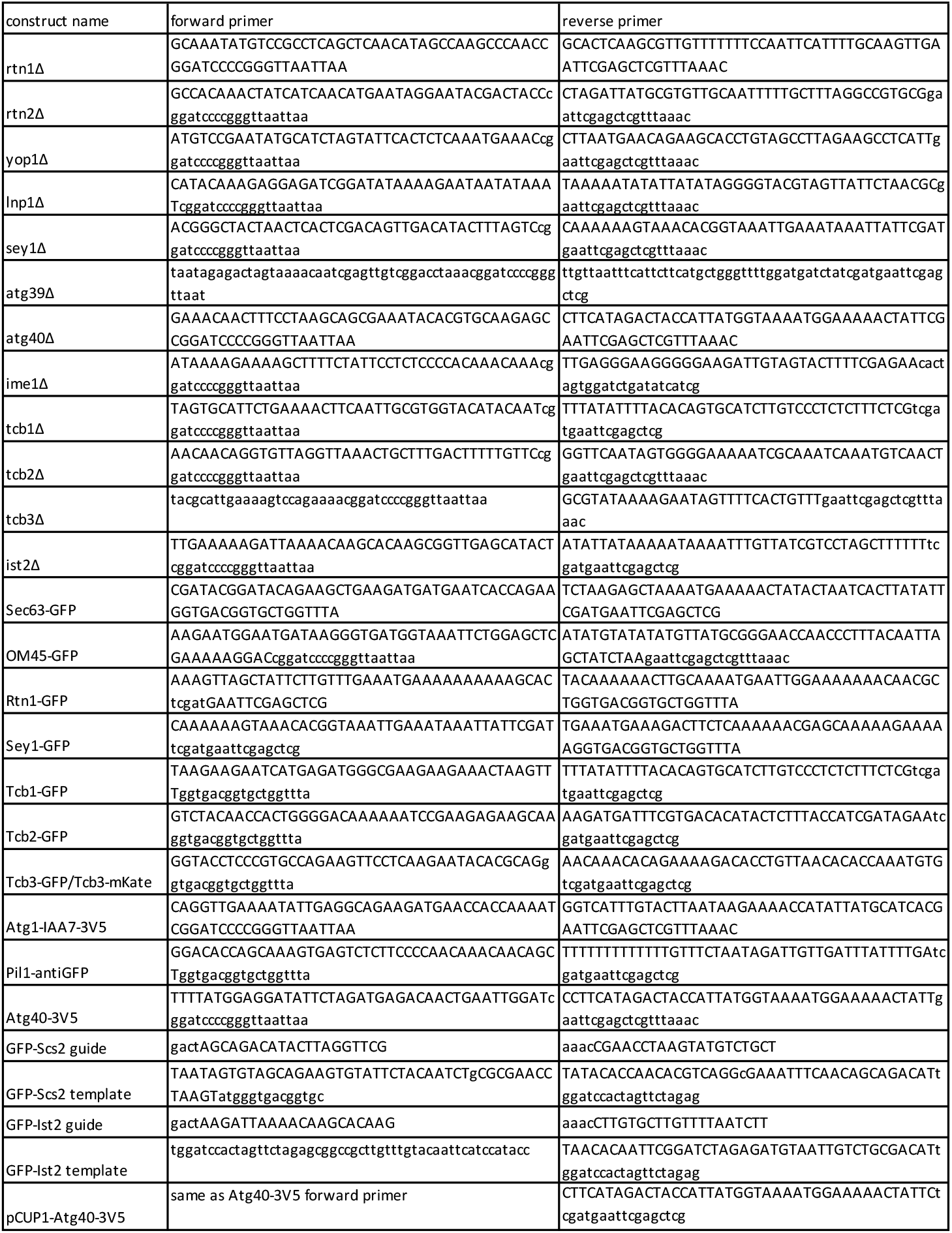
Primers used in this study.

**Table S3.** Plasmids used in this study.

| plasmid name | description |
| --- | --- |
| pÜB1 | pFA6A-KanMX |
| pÜB153 | pFA6A-NatMX |
| pÜB182 | pFA6A-eGFP-KanMX |
| pÜB185 | pFA6A-mKate2-URA3 |
| pÜB217 | pFA6A-hphNT1 |
| pÜB737 | pNH604-pARO10-GFP-HDEL-TRP1 |
| pÜB763 | pFA6A-IAA7-3V5-KanMX |
| pÜB1185 | pNH604-pARO10-mCherry-HDEL-TRP1 |
| pÜB1548 | pRS306-pATG8-linker-eGFP-ATG8 |
| pÜB1707 | pFA6A-link-VH16-caURA3 |
| pÜB1834 | URA/CEN-Cas9-SCS2gRNA |
| pÜB1889 | pRS306-pATG8-linker-eGFP-SCS22 |
| pÜB1977 | pFA6A-pCUP1-ATG40-3V5-hphNT1 |
| pÜB1978 | URA/CEN-Cas9-IST22gRNA |

**Table S4.** Settings used for microscopy experiments.

| Figure 2.x | GFP | RFP | Pol | Z sectioning |
| --- | --- | --- | --- | --- |
| 1A, 1F-G, 2A, 2D, 3A, 3C-E, 4F-G, 5B, S2A-F, S2H-J, S3B-C, S7A | 32% T, 0.025 s | 32% T, 0.025 s | 32% T, 0.1 s | 1 μm, 7 sections |
| 1B, 2C, S2G | 32% T, 0.025 s | 100% T, 0.025 s | 32% T, 0.1 s | 1 μm, 7 sections |
| 2B | 32% T, 0.025 s | 100% T, 0.1 s | 32% T, 0.1 s | 1 μm, 7 sections |
| 2F, 4B-E | 32% T, 0.025 s | 100% T, 0.05 s | 32% T, 0.1 s | 1 μm, 7 sections |
| 6C, 6F, 6I, 7C | 100% T, 0.025 s | 100% T, 0.1 s | 32% T, 0.1 s | 0.5 μm, 15 sections |
| S3A, S3D | 32% T, 0.025 s | 32% T, 0.025 s | 32% T, 0.1 s | 0.5 μm, 15 sections |
| S3E | 100% T, 0.025 s | 32% T, 0.025 s | 32% T, 0.1 s | 0.5 μm, 15 sections |

**Video 1.** Cell depicted in figure 1A.

**Video 2.** Cell depicted in figure S1A.

**Video 3.** Cell depicted in figure 1C.

**Video 4.** Cell depicted in figure 1G.

**Video 5.** Tcb3-GFP cell depicted in figure 2A and S2D.

**Video 6.** Tcb1-GFP cell depicted in figure 2A and S2B.

**Video 7.** Tcb2-GFP cell depicted in figure 2A and S2C.

**Video 8.** GFP-Ist2 cell depicted in figure 2A and S2A.

**Video 9.** GFP-Scs2 cell depicted in figure 2A and S2E.

**Video 10.** GFP-Scs22 cell depicted in figure 2A and S2F.

**Video 11.** Cell depicted in figure 2D.

**Video 12.** Cell depicted in figure S2G.

**Video 13.** Cell depicted in figure S2J.

**Video 14.** Cell depicted in figure S2H.

**Video 15.** Cell depicted in figure S2I.

**Video 16.** Cell depicted in figure 2G.

**Video 17.** Cell depicted in figure 3A.

**Video 18.** Cell depicted in figure S3D.

**Video 19.** Cell depicted in figure 3C.

**Video 20.** Cell depicted in figure 3E.

**Video 21.** Cell depicted in figure 4B.

**Video 22.** Cell depicted in figure 4C.

**Video 23.** Cell depicted in figure 4D.

**Video 24.** Cell depicted in figure 4E.

**Video 25.** Cell depicted in figure 4F.

**Video 26.** Cell depicted in figure 4G.

**Video 27.** Cell depicted in figure 5B.

**Video 28.** Cell depicted in figure S6.

## References

An, H., Ordureau, A., Paulo, J.A., Shoemaker, C.J., Denic, V., and Harper, J.W. (2019). TEX264 Is an Endoplasmic Reticulum-Resident ATG8-Interacting Protein Critical for ER Remodeling during Nutrient Stress. Mol. Cell.

Anding, A.L., and Baehrecke, E.H. (2017). Cleaning House: Selective Autophagy of Organelles. Dev. Cell.

Anwar, K., Klemm, R.W., Condon, A., Severin, K.N., Zhang, M., Ghirlando, R., Hu, J., Rapoport, T.A., and Prinz, W.A. (2012). The dynamin-like GTPase Sey1p mediates homotypic ER fusion in S. cerevisiae. J. Cell Biol. 197, 209–217.

Benjamin, K.R., Zhang, C., Shokat, K.M., and Herskowitz, I. (2003). Control of landmark events in meiosis by the CDK Cdc28 and the meiosis-specific kinase Ime2. Genes Dev.

Brar, G. a., Yassour, M., Friedman, N., Regev, A., Ingolia, N.T., Weissman, J.S., a. Brar, G., Yassour, M., Friedman, N., Regev, A., et al. (2012). High-Resolution View of the Yeast Meiotic Program Revealed by Ribosome Profiling. Science (80-.). 335, 552–557.

Carlile, T.M., and Amon, A. (2008). Meiosis I Is Established through Division-Specific Translational Control of a Cyclin. Cell 133, 280–291.

Chen, J., Tresenrider, A., Chia, M., McSwiggen, D.T., Spedale, G., Jorgensen, V., Liao, H., Van Werven, F.J., and Ünal, E. (2017). Kinetochore inactivation by expression of a repressive mRNA. Elife.

Chen, S., Novick, P., and Ferro-Novick, S. (2012). ER network formation requires a balance of the dynamin-like GTPase Sey1p and the Lunapark family member Lnp1p. Nat. Cell Biol. 14, 707–716.

Chen, S., Desai, T., McNew, J.A., Gerard, P., Novick, P.J., and Ferro-Novick, S. (2015). Lunapark stabilizes nascent three-way junctions in the endoplasmic reticulum. Proc. Natl. Acad. Sci. U. S. A.

Chen, S., Cui, Y., Parashar, S., Novick, P.J., and Ferro-Novick, S. (2018). ER-phagy requires Lnp1, a protein that stabilizes rearrangements of the ER network. Proc. Natl. Acad. Sci. U. S. A.

Cheng, Z., Otto, G.M., Powers, E.N., Keskin, A., Mertins, P., Carr, S.A., Jovanovic, M., and Brar, G.A. (2018). Pervasive, Coordinated Protein-Level Changes Driven by Transcript Isoform Switching during Meiosis. Cell 172.

Chu, S., and Herskowitz, I. (1998). Gametogenesis in yeast is regulated by a transcriptional cascade dependent on Ndt80. Mol. Cell.

Clay, L., Caudron, F., Denoth-Lippuner, A., Boettcher, B., Frei, S.B., Snapp, E.L., and Barral, Y. (2014). A sphingolipid-dependent diffusion barrier confines ER stress to the yeast mother cell. Elife.

Cui, Y., Parashar, S., Zahoor, M., Needham, P.G., Mari, M., Zhu, M., Chen, S., Ho, H.C., Reggiori, F., Farhan, H., et al. (2019). A COPII subunit acts with an autophagy receptor to target endoplasmic reticulum for degradation. Science (80-.).

Cunningham, C.N., Williams, J.M., Knupp, J., Arunagiri, A., Arvan, P., and Tsai, B. (2019). Cells Deploy a Two-Pronged Strategy to Rectify Misfolded Proinsulin Aggregates. Mol. Cell.

Eastwood, M.D., Cheung, S.W.T., Lee, K.Y., Moffat, J., and Meneghini, M.D. (2012). Developmentally Programmed Nuclear Destruction during Yeast Gametogenesis. Dev. Cell.

Eisenberg, A.R., Higdon, A., Keskin, A., Hodapp, S., Jovanovic, M., and Brar, G.A. (2018). Precise Post-translational Tuning Occurs for Most Protein Complex Components during Meiosis. Cell Rep.

Espadas, J., Pendin, D., Bocanegra, R., Escalada, A., Misticoni, G., Trevisan, T., Velasco del Olmo, A., Montagna, A., Bova, S., Ibarra, B., et al. (2019). Dynamic constriction and fission of endoplasmic reticulum membranes by reticulon. Nat. Commun.

Estrada, P., Kim, J., Coleman, J., Walker, L., Dunn, B., Takizawa, P., Novick, P., and Ferro-Novick, S. (2003). Myo4p and She3p are required for cortical ER inheritance in Saccharomyces cerevisiae. J. Cell Biol. 163, 1255–1266.

Forrester, A., De Leonibus, C., Grumati, P., Fasana, E., Piemontese, M., Staiano, L., Fregno, I., Raimondi, A., Marazza, A., Bruno, G., et al. (2019). A selective ER-phagy exerts procollagen quality control via a Calnexin-FAM 134B complex. EMBO J.

Fumagalli, F., Noack, J., Bergmann, T.J., Presmanes, E.C., Pisoni, G.B., Fasana, E., Fregno, I., Galli, C., Loi, M., Soldà, T., et al. (2016). Translocon component Sec62 acts in endoplasmic reticulum turnover during stress recovery. Nat. Cell Biol. 18, 1173–1184.

Gibson, D.G., Young, L., Chuang, R.Y., Venter, J.C., Hutchison, C.A., and Smith, H.O. (2009). Enzymatic assembly of DNA molecules up to several hundred kilobases. Nat. Methods.

Goodman, J.S., King, G.A., and Ünal, E. (2020). Cellular quality control during gametogenesis. Exp. Cell Res.

Graef, M., Friedman, J.R., Graham, C., Babu, M., and Nunnari, J. (2013). ER exit sites are physical and functional core autophagosome biogenesis components. Mol. Biol. Cell.

Guo, Y., Li, D.D., Zhang, S., Yang, Y., Liu, J.J., Wang, X., Liu, C., Milkie, D.E., Moore, R.P., Tulu, U.S., et al. (2018). Visualizing Intracellular Organelle and Cytoskeletal Interactions at Nanoscale Resolution on Millisecond Timescales. Cell 175, 1430–1442.e17.

Hu, J., Shibata, Y., Voss, C., Shemesh, T., Li, Z., Coughlin, M., Kozlov, M.M., Rapoport, T.A., and Prinz, W.A. (2008). Membrane proteins of the endoplasmic reticulum induce high-curvature tubules. Science (80-.).

Hu, J., Shibata, Y., Zhu, P.P., Voss, C., Rismanchi, N., Prinz, W.A., Rapoport, T.A., and Blackstone, C. (2009). A Class of Dynamin-like GTPases Involved in the Generation of the Tubular ER Network. Cell 138, 549–561.

Janke, C., Magiera, M.M., Rathfelder, N., Taxis, C., Reber, S., Maekawa, H., Moreno-Borchart, A., Doenges, G., Schwob, E., Schiebel, E., et al. (2004). A versatile toolbox for PCR-based tagging of yeast genes: New fluorescent proteins, more markers and promoter substitution cassettes. Yeast.

Khaminets, A., Heinrich, T., Mari, M., Grumati, P., Huebner, A.K., Akutsu, M., Liebmann, L., Stolz, A., Nietzsche, S., Koch, N., et al. (2015). Regulation of endoplasmic reticulum turnover by selective autophagy. Nature.

King, G.A., Goodman, J.S., Schick, J.G., Chetlapalli, K., Jorgens, D.M., McDonald, K.L., and Ünal, E. (2019). Meiotic cellular rejuvenation is coupled to nuclear remodeling in budding yeast. Elife.

Lee, B.H., and Amon, A. (2003). Role of Polo-like kinase CDC5 in programming meiosis I chromosome segregation. Science (80-.).

Longtine, M.S., McKenzie, A., Demarini, D.J., Shah, N.G., Wach, A., Brachat, A., Philippsen, P., and Pringle, J.R. (1998). Additional modules for versatile and economical PCR-based gene deletion and modification in Saccharomyces cerevisiae. Yeast.

Manford, A.G., Stefan, C.J., Yuan, H.L., MacGurn, J.A., and Emr, S.D. (2012). ER-to-Plasma Membrane Tethering Proteins Regulate Cell Signaling and ER Morphology. Dev. Cell 23.

Matos, J., Lipp, J.J., Bogdanova, A., Guillot, S., Okaz, E., Junqueira, M., Shevchenko, A., and Zachariae, W. (2008). Dbf4-Dependent Cdc7 Kinase Links DNA Replication to the Segregation of Homologous Chromosomes in Meiosis I. Cell.

Mochida, K., Oikawa, Y., Kimura, Y., Kirisako, H., Hirano, H., Ohsumi, Y., and Nakatogawa, H. (2015). Receptor-mediated selective autophagy degrades the endoplasmic reticulum and the nucleus. Nature.

Morishita, H., and Mizushima, N. (2019). Diverse cellular roles of autophagy. Annu. Rev. Cell Dev. Biol.

Nakanishi, H., De Los Santos, P., and Neiman, A.M. (2004). Positive and Negative Regulation of a SNARE Protein by Control of Intracellular Localization. Mol. Biol. Cell.

Neiman, A.M. (2011). Sporulation in the budding yeast Saccharomyces cerevisiae. Genetics 189, 737–765.

Nishimura, K., Fukagawa, T., Takisawa, H., Kakimoto, T., and Kanemaki, M. (2009). An auxin-based degron system for the rapid depletion of proteins in nonplant cells. Nat. Methods.

Okada, M., Kusunoki, S., Ishibashi, Y., and Kito, K. (2017). Proteomics analysis for asymmetric inheritance of preexisting proteins between mother and daughter cells in budding yeast. Genes to Cells.

Orso, G., Pendin, D., Liu, S., Tosetto, J., Moss, T.J., Faust, J.E., Micaroni, M., Egorova, A., Martinuzzi, A., a McNew, J., et al. (2009). Homotypic fusion of ER membranes requires the dynamin-like GTPase Atlastin. Nature 460, 978–983.

Öztürk, Z., O’Kane, C.J., and Pérez-Moreno, J.J. (2020). Axonal Endoplasmic Reticulum Dynamics and Its Roles in Neurodegeneration. Front. Neurosci.

Petkovic, M., Jemaiel, A., Daste, F., Specht, C.G., Izeddin, I., Vorkel, D., Verbavatz, J.M., Darzacq, X., Triller, A., Pfenninger, K.H., et al. (2014). The SNARE Sec22b has a non-fusogenic function in plasma membrane expansion. Nat. Cell Biol.

Piña, F.J., and Niwa, M. (2015). The ER Stress Surveillance (ERSU) pathway regulates daughter cell ER protein aggregate inheritance. Elife.

Powers, R.E., Wang, S., Liu, T.Y., and Rapoport, T.A. (2017). Reconstitution of the tubular endoplasmic reticulum network with purified components. Nature.

Renvoisé, B., and Blackstone, C. (2010). Emerging themes of ER organization in the development and maintenance of axons. Curr. Opin. Neurobiol. 20, 531–537.

Rossanese, O.W., Reinke, C.A., Bevis, B.J., Hammond, A.T., Sears, I.B., O’Connor, J., and Glick, B.S. (2001). A role for actin, Cdc1 p, and Myo2p in the inheritance of late Golgi elements in Saccharomyces cerevisiae. J. Cell Biol.

Rouskin, S., Zubradt, M., Washietl, S., Kellis, M., and Weissman, J.S. (2014). Genomewide probing of RNA structure reveals active unfolding of mRNA structures in vivo. Nature.

Sawyer, E.M., Joshi, P.R., Jorgensen, V., Yunus, J., Berchowitz, L.E., and Ünal, E. (2019). Developmental regulation of an organelle tether coordinates mitochondrial remodeling in meiosis. J. Cell Biol.

Schmit, H.L., Kraft, L.M., Lee-Smith, C.F., and Lackner, L.L. (2018). The role of mitochondria in anchoring dynein to the cell cortex extends beyond clustering the anchor protein. Cell Cycle.

Schuck, S., Prinz, W.A., Thorn, K.S., Voss, C., and Walter, P. (2009). Membrane expansion alleviates endoplasmic reticulum stress independently of the unfolded protein response. J. Cell Biol. 187, 525–536.

Schwarz, D.S., and Blower, M.D. (2016). The endoplasmic reticulum: Structure, function and response to cellular signaling. Cell. Mol. Life Sci. 73, 79–94.

Shcheprova, Z., Baldi, S., Frei, S.B., Gonnet, G., and Barral, Y. (2008). A mechanism for asymmetric segregation of age during yeast budding. Nature.

Suda, Y., Nakanishi, H., Mathieson, E.M., and Neiman, A.M. (2007). Alternative modes of organellar segregation during sporulation in Saccharomyces cerevisiae. Eukaryot. Cell 6, 2009–2017.

Sugiyama, S., and Tanaka, M. (2019). Distinct segregation patterns of yeast cell-peripheral proteins uncovered by a method for protein segregatome analysis. Proc. Natl. Acad. Sci. U. S. A.

Takizawa, P.A., DeRisi, J.L., Wilhelm, J.E., and Vale, R.D. (2000). Plasma membrane compartmentalization in yeast by messenger RNA transport and a septin diffusion barrier. Science (80-.).

Topolska, M., Roelants, F.M., Si, E.P., and Thorner, J. (2020). TORC2-Dependent Ypk1-Mediated Phosphorylation of Lam2/Ltc4 Disrupts Its Association with the β-Propeller Protein Laf1 at Endoplasmic Reticulum-Plasma Membrane Contact Sites in the Yeast Saccharomyces cerevisiae. Biomolecules.

Ünal, E., Kinde, B., and Amon, A. (2011). Gametogenesis eliminates age-induced cellular damage and resets life span in yeast. Science (80-.).

Voeltz, G.K., Prinz, W.A., Shibata, Y., Rist, J.M., and Rapoport, T.A. (2006). A class of membrane proteins shaping the tubular endoplasmic reticulum. Cell 124, 573–586.

Walter, P., and Ron, D. (2011). The unfolded protein response: From stress pathway to homeostatic regulation. Science (80-.).

Wang, S., Tukachinsky, H., Romano, F.B., and Rapoport, T.A. (2016). Cooperation of the ER-shaping proteins atlastin, lunapark, and reticulons to generate a tubular membrane network. Elife 5.

Wen, F.P., Guo, Y.S., Hu, Y., Liu, W.X., Wang, Q., Wang, Y.T., Yu, H.Y., Tang, C.M., Yang, J., Zhou, T., et al. (2016). Distinct temporal requirements for autophagy and the proteasome in yeast meiosis. Autophagy.

Westrate, L.M.M., Lee, J.E.E., Prinz, W.A.A., and Voeltz, G.K.K. (2015). Form Follows Function: The Importance of Endoplasmic Reticulum Shape. Annu. Rev. Biochem. 84, 791–811.

Wittebolle, L., Marzorati, M., Clement, L., Balloi, A., Daffonchio, D., Heylen, K., De Vos, P., Verstraete, W., and Boon, N. (2009). Initial community evenness favours functionality under selective stress. Nature.

Xu, L., Ajimura, M., Padmore, R., Klein, C., and Kleckner, N. (1995). NDT80, a meiosis-specific gene required for exit from pachytene in Saccharomyces cerevisiae. Mol. Cell. Biol.

Zhang, X., Ding, X., Marshall, R.S., Paez-Valencia, J., Lacey, P., Vierstra, R.D., and Otegui, M.S. (2020). Reticulon proteins modulate autophagy of the endoplasmic reticulum in maize endosperm. Elife.

